# Evolutionary epidemiology of *Streptococcus iniae*: linking mutation rate dynamics with adaptation to novel immunological landscapes

**DOI:** 10.1101/355412

**Authors:** Oleksandra Silayeva, Jan Engelstädter, Andrew C Barnes

**Affiliations:** The University of Queensland, School of Biological Sciences, St Lucia Campus, Brisbane, Queensland 4072, Australia

**Keywords:** DNA repair deficiency, heritable mutator phenotype, antigenic variation, immune escape, vaccine-induced serotype replacement, host jumps

## Abstract

Pathogens continuously adapt to changing host environments where variation in their virulence and antigenicity is critical to their long-term evolutionary success. The emergence of novel variants is accelerated in microbial mutator strains (mutators) deficient in DNA repair genes, most often from mismatch repair and oxidised-guanine repair systems (MMR and OG respectively). Bacterial MMR/OG mutants are abundant in clinical samples and show increased adaptive potential in experimental infection models, yet the role of mutators in the epidemiology and evolution of infectious disease is not well understood. Here we investigated the role of mutation rate dynamics in the evolution of a broad host range pathogen, *Streptococcus iniae*, using a set of 80 strains isolated globally over 40 years. We have resolved phylogenetic relationships using non-recombinant core genome variants, measured *in vivo* mutation rates by fluctuation analysis, identified variation in major MMR/OG genes and their regulatory regions, and phenotyped the major traits determining virulence in streptococci. We found that both mutation rate and MMR/OG genotype are remarkably conserved within phylogenetic clades but significantly differ between major phylogenetic lineages. Further, variation in MMR/OG loci correlates with occurrence of atypical virulence-associated phenotypes, infection in atypical hosts (mammals), and atypical tissue of a vaccinated primary hosts (barramundi bone). These findings suggest that mutators are likely to facilitate adaptations preceding major diversification events, and may promote emergence of variation permitting colonisation of a novel host tissue, novel host taxa (host jumps), and immune-escape in the vaccinated host.

## 1. Introduction

Microbial evolution during infection can increase pathogen fitness that, while in the host, is largely determined by interactions with immune responses ^1,2^. Variation in virulence and antigenicity determinants allows pathogens to escape immune clearance and spread in a host population ^2,3^. Therefore, antigenic variation plays major role in the epidemiology of infectious diseases and often compromises their control by vaccination ^4,5^. As vertebrate immune responses are remarkably diverse and complex, every individual host and tissue represents a unique ‘immunological niche’ that requires adaptation^6^. In bacteria, host-adaptive variation could be obtained via accrual of mutation, recombination, and transposition ^7^. Notably, genes involved in host-pathogen interactions are often represent “mutation hot-spots” as they contain sequence physically pre-disposed to errors during replication^8^. Further, overall fidelity of replication in bacteria is influenced by the environmental conditions, and elevation of mutation rate, known as stress-induced mutagenesis (SIM), occurs in adapting populations^9,10^. One aspect of SIM is temporary physiological shifts producing higher mutation rate, such as up-regulation of error-prone polymerases that bypass DNA lesions during the bacterial general stress response, SOS ^9,10^. Another aspect of SIM is increased frequency of mutators - heritably hypermutable strains containing mutator alleles, also referred to as constitutive or permanent mutators, to distinguish them from temporary (physiologically) hypermutable cultures^9–11^. Mutator strains emerge via disruption of DNA repair genes, most often belonging to the mismatch-repair (MMR) and oxidized guanine (OG) repair systems ^12,13^. MMR corrects base mis-parings and short insertion/deletion loops that occur during DNA replication ^14,15^. It also prevents incorporation of divergent DNA sequences via non-homologous and homeologous (partially homologous) recombination; thus MMR disfunction produces not only hypermutable but also hyper-recombinable phenotypes^15,16^. The OG system repairs oxidative DNA damage throughout the cell cycle^17,18^. Although relatively very conserved ^19^, MMR and OG genes are quite variable among major bacterial pathogens implicating mutation rate dynamics in the evolution of virulence and antigenicity^20,21^.

In adapting populations, mutator alleles are indirectly selected (“hitchhike”) along with stress-specific resistance variants, such as antibiotic resistance or host-immunity resistance mutations^22–24^. Considering pathogens adapt to every individual host, the host population represents an extremely heterogeneous environment where adaptive processes are ongoing and mutator alleles are likely to persist^25^. Indeed, bacterial mutators are abundant among clinical isolates, especially in chronic cases^11,26^. Although it may be partly attributable to antibiotic selection^27–29^, the association between frequency of mutators in diagnostic samples and antibiotic treatments is weak^30,31^. Further, unexposed to antimicrobials, knockout mutator strains show shifts in virulence ^32,33^ as well as increased colonization potential in mice^34–36^. Thus, prevalence of mutators in clinical isolates most likely results from selective process for host-adaptive mutations during infection ^30,32,37,38^.

Although mutator alleles can accelerate short-term adaptation they are also indirectly selected against along with deleterious variants^24,39–41^ , and emerging anti-mutator genotypes start to dominate in the adapted mutator clone^42,43^. When copy-number variant confers a mutator phenotype the reverse mutation can fully restore the non-mutator genotype and phenotype^44^. The latter is also possible when prophage integration confers the mutator phenotype and excision the non-mutator phenotype or vice versa^45^. However, when anti-mutator alleles originate via random local mutation or recombination in a mutator strain, the final adapted mutant is likely to have a mutation rate slightly different to the original mutation rate along with traceable shifts MMR/OG genotype^43,46,47^. Therefore, we hypothesized that mutation rate dynamics produced by emergence of mutator and compensatory anti-mutator alleles could be investigated and linked to adaptive evolution via coupling of comparative phylogenomics with mutation rate and pathogenicity phenomics. Here we used *Streptococcus iniae*, a rapidly evolving aquatic pathogen with global distribution, broad-host range, and record of re-infection of previously immunised farmed fish ^48,49^, as a model organism to evaluate the potential role of mutators in epidemiology. We determined phylogenetic relationship based on non-recombinant core genome single nucleotide polymorphisms (SNPs), identified variation in the *in vivo* mutation rates estimated by fluctuation analysis ^50,51^, and variation in major MMR (*mutS*, *mutL*, *dnaN*, *recD2*, *rnhC*) ^52^ and OG (*mutY*, *mutM*, *mutT*)^21^ coding and regulatory loci among 80 diverse strains of *S. iniae* isolated globally over 40 years. This revealed that shifts in mutation rate and MMR/OG genotype correlate with phylogenetic diversifications of *S. iniae*, but, in contrast, are highly conserved within phylogenetic clades. To determine whether diversification is mainly driven by the host-adaptive evolution, we identify variation in major phenotypic traits that determine virulence in streptococci (capsular polysaccharide, hemolysin, length of cell chains) ^53–55^ and bacteria in general (resistance to reactive oxygen species (ROS), and biofilm formation) ^56,57^. We find that occurrence of atypical phenotypes for these traits correlates strongly with number of SNPs in DNA repair loci, and prevails in isolates from unusual immunological landscapes such as mammalian host and bone tissue of vaccinated fish. This suggests that fluctuations in mutation rate may promote evolution of virulence, transmission between divergent host species, and occurrence of vaccine-escape mutants.

## 2. Materials and methods

### Strains and growth conditions

Eighty isolates of *S. iniae* collected in Australia, USA, Canada, Israel, Honduras, and Thailand between 1976 and 2016 from eight fish species (*Lates calcarifer*, *Scortum barcoo, Epalzeorhynchos frenatum, Epalzeorhynchos bicolor, Oreochromis sp., Channa striata, Chromobotia macracanthus, Oncorhynchus mykiss*) and three mammalian species (*Homo sapiens, Inia geoffrensis, Pteropus alecto*) were analysed (Table 1). Strains were received from culture collections, veterinarians, or directly from fish farms (Suppl. Table 3) and stored as master seed stocks without further subculture at −80°C in Todd-Hewitt Broth (THB, Oxoid) + 20% glycerol. Frozen stocks were recovered on Columbia agar supplemented with 5% defibrinated sheep blood (Oxoid), and cultured at 28°C on Todd-Hewitt agar (THA) or in THB with agitation 200 rpm unless otherwise specified.

**Table 1.** *Streptococcus iniae* strains used in this study. Includes origin details (host species, time, site of isolation), phylogenetic affiliation, virulence-associated phenotypes, and mutation rate. Atypical places of isolation (hosts, tissues) and deviant phenotypes are in bold.

| Strain | Clade | Host | Year | Location | Capsule | Hemolysis | Chains | Oxidation resistance | Biofilm formation | Mutation rate (10<sup>-7</sup>) |
| --- | --- | --- | --- | --- | --- | --- | --- | --- | --- | --- |
| QMA0071 | A | *Lates calcarifer* | 2000 | QLD | wildtype | wildtype | wildtype | wildtype | wildtype | 0.2041 |
| QMA0074 | C2 | *Lates calcarifer* | 1998 | QLD | **absent** | wildtype | **long** | wildtype | wildtype | 0.6579 |
| QMA0077 | C2 | *Lates calcarifer* | 1995 | QLD | **absent** | wildtype | **long** | wildtype | wildtype | 0.7067 |
| QMA0078 | A | *Lates calcarifer* | 2001 | QLD | wildtype | wildtype | wildtype | wildtype | wildtype | 0.2153 |
| QMA0080 | C1 | *Lates calcarifer* | 2004 | WA | wildtype | wildtype | wildtype | wildtype | wildtype | 0.1966 |
| QMA0082 | C1 | *Lates calcarifer* | 2004 | WA | wildtype | wildtype | wildtype | wildtype | wildtype | 0.1761 |
| QMA0083 | A | *Lates calcarifer* | 2004 | WA | wildtype | wildtype | wildtype | wildtype | wildtype | 0.1894 |
| QMA0084 | B | *Epalzeorhynchus kalopterus* | 2001 | WA | wildtype | **weak** | wildtype | wildtype | wildtype | 0.7293 |
| QMA0087 | A | *Lates calcarifer* | 2004 | WA | wildtype | wildtype | wildtype | wildtype | wildtype | 0.1925 |
| QMA0130 | E2 | ***Homo sapiens*** | 1995 | Canada | **absent** | wildtype | wildtype | wildtype | wildtype | 0.468 |
| QMA0131 | E2 | ***Homo sapiens*** | 1995 | Canada | **absent** | wildtype | wildtype | wildtype | wildtype | 0.4488 |
| QMA0133 | E1 | ***Homo sapiens*** | 2001 | USA | **absent** | wildtype | wildtype | wildtype | **increased** | 0.5539 |
| QMA0134 | E1 | ***Homo sapiens*** | 2001 | USA | **differential** | wildtype | wildtype | wildtype | **increased** | 0.5457 |
| QMA0135 | E1 | ***Homo sapiens*** | 2002 | USA | wildtype | wildtype | **short** | **increased** | wildtype | 0.631 |
| QMA0137 | E1 | ***Homo sapiens*** | 2004 | USA | wildtype | wildtype | **short** | **increased** | wildtype | 0.5495 |
| QMA0138 | E1 | ***Homo sapiens*** | 2004 | USA | wildtype | wildtype | **short** | **increased** | wildtype | 0.5989 |
| QMA0139 | F | *Fish sp.* | 1996 | Canada | **absent** | wildtype | wildtype | wildtype | wildtype | 0.4311 |
| QMA0140 | - | ***Inia geoffrensis*** | 1976 | USA | **absent** | wildtype | wildtype | **increased** | **increased** | 0.0901 |
| QMA0141 | - | ***Inia geoffrensis*** | 1978 | USA | **absent** | wildtype | wildtype | **increased** | **increased** | 1.0884 |
| QMA0142 | C1 | *Lates calcarifer* | 2005 | NT | wildtype | wildtype | wildtype | wildtype | wildtype | 0.2044 |
| QMA0150 | C1 | *Lates calcarifer* | 2005 | NT | wildtype | wildtype | wildtype | wildtype | wildtype | 0.1952 |
| QMA0155 | A | *Lates calcarifer* | 2005 | NSW RAS | wildtype | wildtype | wildtype | wildtype | wildtype | 0.1493 |
| QMA0156 | A | *Lates calcarifer* | 2005 | NSW RAS | wildtype | wildtype | wildtype | wildtype | wildtype | 0.1666 |
| QMA0157 | A | *Lates calcarifer* | 2005 | NSW RAS | wildtype | wildtype | wildtype | wildtype | wildtype | 0.1545 |
| QMA0158 | A | *Lates calcarifer* | 2006 | SA RAS | **absent** | wildtype | **long** | wildtype | wildtype | 0.1678 |
| QMA0159 | A | *Lates calcarifer* | 2006 | SA RAS | wildtype | wildtype | wildtype | wildtype | wildtype | 0.2054 |
| QMA0160 | A | *Lates calcarifer* | 1999 | SA RAS | wildtype | wildtype | wildtype | wildtype | wildtype | 0.1481 |
| QMA0161 | A | *Lates calcarifer* | 2000 | SA RAS | wildtype | wildtype | wildtype | wildtype | wildtype | 0.1742 |
| QMA0162 | A | *Lates calcarifer* | 2000 | SA RAS | wildtype | wildtype | wildtype | wildtype | wildtype | 0.1718 |
| QMA0163 | A | *Lates calcarifer* | 2000 | SA RAS | wildtype | wildtype | wildtype | wildtype | wildtype | 0.1523 |
| QMA0164 | C2 | *Lates calcarifer* | 2006 | QLD | wildtype | wildtype | wildtype | wildtype | wildtype | 0.7102 |
| QMA0165 | C2 | *Lates calcarifer* | 2006 | QLD | wildtype | wildtype | wildtype | wildtype | wildtype | 0.6606 |
| QMA0177 | C1 | *Lates calcarifer* | 2006 | NT | wildtype | wildtype | wildtype | wildtype | wildtype | 0.1718 |
| QMA0180 | C1 | *Lates calcarifer* | 2006 | NT | wildtype | wildtype | wildtype | wildtype | wildtype | 0.2011 |
| QMA0186 | D | *Oncorhynchus mykiss* | 2000 | Israel | wildtype | **weak** | wildtype | wildtype | **increased** | 0.5135 |
| QMA0187 | - | *Channa striata* | 1983 | Thailand | **absent** | **weak** | wildtype | wildtype | wildtype | 0.4635 |
| QMA0188 | D | *Oncorhynchus mykiss* | 1998 | Israel | wildtype | **weak** | wildtype | wildtype | **increased** | 0.5042 |
| QMA0189 | D | *Oncorhynchus mykiss* | 1996 | Reunion | wildtype | **weak** | wildtype | wildtype | wildtype | 0.4907 |
| QMA0190 | F | *Channa striata* | 1988 | Thailand | **absent** | wildtype | wildtype | wildtype | wildtype | 0.6413 |
| QMA0191 | C1 | *Lates calcarifer* | 2005 | NT | wildtype | wildtype | wildtype | wildtype | wildtype | 0.1883 |
| QMA0207 | C1 | *Lates calcarifer* | 2006 | NT | wildtype | wildtype | wildtype | wildtype | wildtype | 0.1486 |
| QMA0216 | A | *Lates calcarifer* | 2007 | QLD | **absent** | wildtype | **long** | wildtype | wildtype | 0.1949 |
| QMA0218 | C2 | *Lates calcarifer* | 2007 | QLD | wildtype | wildtype | wildtype | wildtype | wildtype | 0.7093 |
| QMA0220 | A | *Lates calcarifer* | 2006 | NSW RAS | wildtype | wildtype | wildtype | wildtype | wildtype | 0.2086 |
| QMA0221 | A | *Lates calcarifer* | 2007 | NSW RAS | wildtype | wildtype | wildtype | wildtype | wildtype | 0.1814 |
| QMA0222 | A | *Lates calcarifer* | 2006 | SA RAS | wildtype | wildtype | wildtype | wildtype | wildtype | 0.1829 |
| QMA0233 | F | *Lates calcarifer, bone* | 2009 | NSW RAS | **absent** | **weak** | **long** | wildtype | **increased** | 0.362 |
| QMA0234 | F | *Lates calcarifer, bone* | 2009 | NSW RAS | **absent** | **weak** | **long** | wildtype | **increased** | 0.354 |
| QMA0235 | F | *Lates calcarifer, bone* | 2009 | NSW RAS | **absent** | **weak** | **long** | wildtype | **increased** | 0.3496 |
| QMA0236 | F | *Lates calcarifer, bone* | 2009 | NSW RAS | **absent** | **weak** | **long** | wildtype | **increased** | 0.3413 |
| QMA0244 | A | *Lates calcarifer* | 2008 | SA RAS | wildtype | wildtype | wildtype | wildtype | wildtype | 0.2101 |
| QMA0245 | A | *Lates calcarifer* | 2008 | SA RAS | wildtype | wildtype | wildtype | wildtype | wildtype | 0.1988 |
| QMA0246 | A | *Lates calcarifer* | 2009 | SA RAS | wildtype | wildtype | wildtype | wildtype | wildtype | 0.2149 |
| QMA0247 | A | *Lates calcarifer* | 2009 | SA RAS | wildtype | wildtype | wildtype | wildtype | wildtype | 0.2059 |
| QMA0248 | A | *Lates calcarifer* | 2009 | SA RAS | wildtype | wildtype | wildtype | wildtype | wildtype | 0.1893 |
| QMA0249 | F | *Lates calcarifer, bone* | 2009 | SA RAS | **differential** | **weak** | **long** | wildtype | **increased** | 0.3817 |
| QMA0250 | A | *Lates calcarifer* | 2007 | NSW RAS | wildtype | wildtype | wildtype | wildtype | wildtype | 0.195 |
| QMA0251 | A | *Lates calcarifer* | 2008 | NSW RAS | wildtype | wildtype | wildtype | wildtype | wildtype | 0.1927 |
| QMA0252 | A | *Lates calcarifer* | 2008 | NSW RAS | wildtype | wildtype | wildtype | wildtype | wildtype | 0.2006 |
| QMA0253 | F | *Lates calcarifer, bone* | 2009 | NSW RAS | **absent** | wildtype | **long** | wildtype | **increased** | 0.3555 |
| QMA0254 | F | *Lates calcarifer, bone* | 2009 | NSW RAS | **absent** | wildtype | **long** | wildtype | **increased** | 0.3712 |
| QMA0258 | A | *Lates calcarifer* | 2008 | QLD | wildtype | wildtype | wildtype | wildtype | wildtype | 0.1876 |
| QMA0371 | A | *Scortum barcoo* | 2011 | NSW | wildtype | wildtype | wildtype | wildtype | wildtype | 0.1889 |
| QMA0373 | C2 | *Lates calcarifer* | 2012 | QLD | wildtype | wildtype | wildtype | wildtype | wildtype | 0.6908 |
| QMA0374 | C2 | *Lates calcarifer* | 2012 | QLD | wildtype | wildtype | wildtype | wildtype | wildtype | 0.729 |
| QMA0445 | G | *Oreochromis sp.* | 1998 | USA | **absent** | wildtype | wildtype | wildtype | wildtype | 0.2149 |
| QMA0446 | G | *Oreochromis sp.* | 1998 | USA | **absent** | wildtype | wildtype | wildtype | wildtype | 0.1687 |
| QMA0447 | E1 | *Hybrid striped bass* | 1996 | USA | wildtype | wildtype | **short** | **increased** | wildtype | 0.5902 |
| QMA0448 | E1 | *Hybrid striped bass* | 1998 | USA | wildtype | wildtype | **short** | **increased** | wildtype | 0.5529 |
| QMA0457 | A | *Oreochromis sp.* | 2005 | USA | wildtype | wildtype | wildtype | wildtype | wildtype | 0.1682 |
| QMA0458 | A | *Epalzeorhynchus bicolor* | 2004 | USA | wildtype | wildtype | wildtype | wildtype | wildtype | 0.1778 |
| QMA0462 | B | *Chromobotia macracanthus* | 2005 | USA | wildtype | **weak** | wildtype | wildtype | wildtype | 0.4242 |
| QMA0463 | B | *Chromobotia macracanthus* | 2005 | USA | wildtype | **weak** | wildtype | wildtype | wildtype | 0.414 |
| QMA0466 | E2 | *Oreochromis sp.* | - | USA | wildtype | wildtype | **short** | wildtype | wildtype | 0.4602 |
| QMA0467 | A | *Epalzeothynchos frenatum* | 2004 | USA | wildtype | wildtype | wildtype | wildtype | wildtype | 0.1695 |
| QMA0468 | A | *Oreochromis sp.* | 2005 | USA | wildtype | wildtype | wildtype | wildtype | wildtype | 0.1689 |
| QMA0490 | A | *Oreochromis sp.* | 2015 | Honduras | wildtype | wildtype | wildtype | wildtype | wildtype | 0.1812 |
| QMA0491 | A | *Oreochromis sp.* | 2015 | Honduras | wildtype | wildtype | wildtype | wildtype | wildtype | 0.2 |
| QMA0492 | A | *Oreochromis sp.* | 2015 | Honduras | wildtype | wildtype | wildtype | wildtype | wildtype | 0.1849 |
| QMA0493 | A | *Oreochromis sp.* | 2016 | Honduras | wildtype | wildtype | wildtype | wildtype | wildtype | 0.1987 |

### Estimation of mutation rates by fluctuation analysis

A fluctuation analysis assay for spontaneous occurrence of rifampicin resistance was optimized according to Rosche and Foster ^58^. A single broth culture was initiated from five separate colonies, recovered on Columbia blood agar from stock, and grown overnight to late-exponential phase in THB. Cultures were adjusted to OD_600_ = 1 (10^8^ CFU/mL), diluted 1:100, and distributed in 200 μl aliquots into 8 wells of a sterile U-bottom 96-well plate (Greiner). Prior to dilution, 100 μl of each OD_600_-adjusted culture was spread onto THA containing 0.5 μg/mL rifampicin to confirm absence of resistant mutants. Although comparatively large initial inocula (1-2 × 10^5^ CFU per culture, confirmed by Miles and Misra CFU count ^59^ of OD-adjusted cultures) were used, this minimised variance in final CFU number (N_t_) determined in preliminary experiments. Invariability in N_t_ allowed statistical comparison of mutation rates by the Maximum Likelihood Ratio test ^60^. To infer the N_t_ in final cultures and monitor their growth by optical density, replicate plates containing two cultures per strain were prepared and incubated in a BMG FLUOstar OPTIMA microplate reader. When these representative cultures entered early stationary phase, CFU counts were performed by Miles and Misra method ^59^ and invariably estimated as 1-2 × 10^8^ CFU in each well. Immediately after, entire 200 μL cultures (above) were plated on THA containing 0.5 μg/mL rifampicin for selection of mutants, dried under laminar flow, and incubated until rifampicin resistant colonies appeared. The assay was repeated four times for each strain in blocks of eight cultures using 20 isolates haphazardly chosen from different phylogenetic lineages in each measurement. In some cases, e.g. a single strain representing an independent phylogenetic lineage or with significantly differences in mutation rates detected in closely related strains, assays were repeated 1-2 more times to increase statistical power. The rifampicin-resistant mutant counts were pooled into single data sets representing 32 to 48 cultures per strain (Suppl. Table 2).

### DNA extraction, preparation and sequencing

Genomic DNA was extracted from cells collected from 10 mL late-exponential phase culture in Todd-Hewitt broth with the DNeasy Blood & Tissue kit (Qiagen) using a modified protocol with an additional lysis step as described previously ^61^. Sequencing was performed on the Illumina HiSeq2000 platform from Nextera XT pair-end libraries at Australian Genome Research Facility, Melbourne. A reference genome from strain QMA0248 was constructed using both long reads derived from a single Smrt Cell using the PacBio RS II system with P4C2 chemistry and short reads from Illumina HiSeq2000 derived from Nextera XT paired-end libraries as reported elsewhere (NCBI accession no: GCA_002220115.1). All sequence data are deposited at NCBI under Bioproject number PRJNA417543, SRA accession SRP145425. Sample numbers, accession numbers and extended metadata are provided in Suppl. Table 3. Assembly statistics are provided in Suppl. Table 4.

### Genome assembly, recombination detection and phylogenetic analysis

Phylogeny was constructed based on core genome SNPs from *de novo* genome assemblies filtered to remove recombination breakpoints. Paired-end reads from Illumina were trimmed with Nesoni clip tool version 0.132 (http://www.vicbioinformatics.com/software.nesoni.shtml), with minimum read length 50, and the first 15 bp of each read removed as quality deterioration in this region was observed when assessed with FASTQC version 0.11.5. Assembly was performed using the SPAdes assembler version 3.7.1 ^62^, with minimum read coverage cutoff set to 10. Quality of assemblies was assessed with QUAST 3.2 ^63^. Contigs were ordered by alignment to QMA248 reference genome (CP022392.1) with Mauve Contig Mover 2.4.0 ^64^. Genome annotation was performed using Prokka 1.11 ^65^. Rapid alignment of core genomes was carried out using parsnp in the Harvest Tools suite version 1.2 ^66^, and the resulting alignment provided as an input to Gubbins 1.4.7 ^67^ for detection and exclusion of variants produced by recombination. Phylogenies were then inferred from post-filtered core genome polymorphic sites by maximum likelihood using RAxML 8.2.8 ^68^ with the general time reversible nucleotide substitution model GTRGAMMA and bootstrap support from 1000 iterations. Effect of ascertainment bias associated with using only polymorphic sites on branch length during ML inference was corrected using Felsenstein’s correction implemented in RAxML 8 ^69^. The resulting phylogenetic tree was visualized using Dendroscope v 3.5.7 ^70^ with bootstrap node support value cut-off 75. For the phylogram figure, tip labels were hidden for clarity and the edge containing QMA0141 was re-scaled as dotted line representing 100-fold decrease in length (Fig. 1). A cladogram based on the inferred phylogeny, showing all tip labels and bootstrap support for each node, was annotated with metadata using Evolview V2 (Fig. 2)^71^. To determine whether there was a strong temporal signal in the phylogenetic data, a root-to-tip regression of branch length against time since isolation was performed in TempEst ^72^, using genetic distances corrected for ascertainment bias in an unrooted tree estimated by maximum likelihood from the alignment of non-recombinant core-genome SNPs in RAxML 8.2.8.

### Variation in MMR/OG genes and their regulatory regions, SNP analysis

Large variants in MMR (*dnaN*, mut*S, muTL, recD2, rnhC*) and OG genes (*mutY* , *mutM*, *mutX*) and surrounding regions were identified within multiple alignments of *de novo* genome assemblies using progressiveMauve ^73^. A 56 bp deletion found in the *mutY* promoter of clade B was confirmed by PCR using the CAGAAGGAAGAAACAGAC_F/ACCTCTATTGTAGCAAAG_R primers set. SNP analysis was performed by read-mapping with Geneious version 9.1 ^74^, using default settings unless otherwise specified. Paired-end reads from each genome were trimmed, merged into a single file with expected distances between the reads set to 250 bp, and mapped to the curated reference genome of strain QMA0248. Mapped reads were used to detect SNPs with minimum coverage set to 10 and frequency to 0.9. A consensus “pseudogenome” was generated for every strain based on the reference sequence incorporating any detected variants ^75^. Multiple alignment of these pseudogenomes was carried out with *Geneious* aligning tool to identify SNPs in MMR/OG genes, their ribosome-binding sites, and promoter regions. Promoter regions of repair genes were determined with BPROM ^76^, protein functional domains by SMART genomic ^77^, and effect of amino acid substitutions on protein function by PROVEAN Protein with a sensitivity cut-off of 1.3 ^78^. To estimate the mutational bias among phylogenetic lineages, SNP types and position per strain were exported as a table and SNPs of each type occurring only in each clade but not in any other clade were counted and plotted using a custom script in R (Suppl. Fig. 1).

### Detecting possible association between variation in DNA repair genes and mobilome

A pangenome for the 80 *S. iniae* isolates was constructed with GView server ^79^ using the QMA0248 reference genbank (.gbk) file as seed by sequentially adding 79 genome assemblies by clade in the order derived from the cladogram (Fig. 2). Phage positions in the resulting pangenome were then determined by BLAST using Phaster ^80^ while IS positions were identified using ISFinder ^81^ with a BLAST e-value cutoff of 1 × 10^−30^. Positions of MMR and OG genes in the pangenome were determined via manual search of the .gbk file. An image was created in BRIG ^82^ using the pangenome as reference then aligning the 80 genome assemblies by BLAST with an e-value cutoff of 1 × 10^−30^. The resulting image was annotated in BRIG from a spreadsheet derived from the positions determined above, and coloured according to clade (Suppl. Fig. 2).

### Phenotypic variation related to virulence and antigenicity

A multitude of intracellular and secreted enzymes contribute to virulence in the *Streptococcus* genus ^83,84^. Considering the large number of strains, we limited analysis to major, easily observable, and distinct phenotypes strongly associated with virulence in streptococci (capsule, hemolysis, cell chains) and bacteria in general (oxidation resistance, biofilms). All assays were performed at least in triplicate per strain.

### Buoyant density assay for presence of polysaccharide capsule

Presence/absence of polysaccharide capsule was estimated by Percoll buoyant density assay. Isotonic stock Percoll (ISP) was prepared by mixing nine parts of Percoll with one part of 1.5 M NaCl. Then, 6 parts of ISP was diluted with 4 parts of 0.15 M NaCl to make final 50 % Percoll solution, which was distributed by 3 mL into flow cytometry tubes. THB cultures (10 mL) grown to late-exponential phase were adjusted at OD_600_ 1 (10^8^ CFU/mL), centrifuged at 3220 × *g* for 5 min, resuspended in 0.5 mL of 0.15 M NaCl, and layered onto the Percoll. Tubes were centrifuged at 4°C in a swinging bucket rotor at 4000 × *g* for 3 h with low acceleration and no brake. In this assay, encapsulated cells form a clear compact band in the Percoll gradient (Fig. 3, A1), non-encapsulated cells form a pellet in the bottom of the tube (Figure 3, A2) and, occasionally, strains show differential expression of capsular polysaccharide evidenced by band and a pellet (Fig. 3, A3).

### Haemolytic activity assay on sheep blood agar

Rapid high-throughput detection of impaired haemolytic activity was achieved by blood-agar clearance zone assay. Briefly, 5 mm wells were made in Columbia agar supplemented with 5% defibrinated sheep blood (Oxoid, Australia). Bacterial cultures grown to late-exponential phase (~10^8^ CFU) were diluted 1:1000 and 50 μL of diluted cultures were pipetted into punctures in the agar (initial inoculum ~5×10^4^ CFU in each puncture). A 3 mm wide clearance zone was generally produced by bacterial lysis of sheep erythrocytes during 24 h incubation (Fig. 3, B1). Where haemolysis was absent or fragmentary, impeded haemolytic activity was recorded (Fig. 3, B2).

### Chain formation microscopy

To assess chain formation, *S. iniae* cultures in THB were grown stationarily in 96-well plates at 28 °C for 24 h, mixed and 5 μL wet mounts prepared on a glass microscope slide. Slides were observed by bright field microscopy under 40x objective with an Olympus BX40 microscope and captured using an Olympus DP28 digital camera using CellSens software (Olympus Optical Co, Japan). Specimens were observed for at least 3 min and 2-20 cell chains were generally present (Fig. 3, C1). Where over 20 cells in a chain were repeatedly detected (Fig. 3, C2) increased chain formation was recorded, and when more than 10 cells in a chain were not detected (and the culture was mainly composed of detached cells) (Fig. 3, C3) impeded chain formation was recorded. For the figure, 50 μL of cultures were dried onto slides at RT, fixed with methanol, Gram stained, and images captured under a 100x objective.

### Oxidation resistance assay

Minimum inhibitory concentration (MIC) and minimum bactericidal concentration (MBC) ^85^ of hydrogen peroxide were measured to assess oxidative stress resistance among *S. iniae* strains. THB cultures grown to late exponential phase were adjusted to OD_600_ = 1 (10^8^ CFU/mL), diluted 1000-fold, and distributed by 100 μL (~10^4^ CFU) into wells of a U-bottom 96-well plate (Greiner). THB (100 μL) with 2-fold excess concentration of hydrogen peroxide was added to the wells and serially diluted twofold, resulting in a range from 0 to 10 mM final peroxide concentrations. Plates were incubated stationarily for 24h and examined for presence of observable growth to determine the MIC. Viable cell counts of cultures without visible growth were performed to determine MBC. MIC of hydrogen peroxide was estimated as 3 mM for all strains, and MBC as 4 mM for the majority of strains. MBC elevated to 5 mM in dolphin and human isolates from USA was classified deviation from a wildtype phenotype.

### Biofilm formation and visualisation assay

*S. iniae* biofilms were grown for 4 days in 8-well Lab-Tek® II Chamber Slide™ Systems. To prepare an initial inoculum of 5 × 10^5^ CFU, THB cultures grown to late exponential phase were adjusted to OD_600_ = 1 (10^8^ CFU/mL), diluted 100-fold, and 0.5 mL of diluted cultures were placed into Chamber slide wells. After 24 h incubation, THB was removed and replaced with fresh THB every 10-14 h for 3 days. For visualisation, biofilms were washed in PBS, stained for 15 min with 1 μM fluorescent *Bac*Light Red bacterial stain, washed in PBS, fixed for 30 min with 10% formalin, and washed twice PBS. Z-stacks were collected by ZEISS LSM 710 Inverted Laser Scanning Confocal Microscope at 20x objective and visualised using ZEN2012. Quantification of biofilms was performed using COMSTAT ^86^. Structures of biomass larger than 3.5 and average thickness over 5 were classified as increased biofilm forming activity (Fig. 3 D).

### Identifying wildtype virulence-related phenotypes

We consider prevalent phenotypes as a wildtype, namely: presence of polysaccharide capsule (71.25 % of strains; Figure 3, A1) and haemolytic activity (85% of strains; Figure 3, B1), up to 20 cells in a chain (77.5 % of strains; Figure 3, A1), 3 mM MIC and 4 mM MBC of hydrogen peroxide (91.25 % of strains), and 8 −12 μm thick biofilms covering up to 30% of representative image (86.25 % of strains, Fig. 4A). Other phenotypes are regarded as deviant and discussed.

### Statistical analysis

Analysis of co-variance between number of MMR and OG variants, mutation rates, number of atypical phenotype associated with virulence, and atypical places of isolation were tested with a Phylogenetic Generalised Least Squares model that accounts for relatedness among strains under Pagel’s λ of 0.6 implemented in R ^87^. Estimation of mutation rates was carried out using the Ma-Sandri-Sarkar Maximum Likelihood Estimation (MSS-MLE) method as implemented in FALCOR fluctuation analysis calculator ^50^. Differences in mutation rates were compared by Likelihood Ratio Test (LRT) using the rSalvador R package ^60,88^. LRT estimates the overlap between confidence intervals calculated for two fluctuation experiment data sets and allows to compare fluctuation assay data with different number of cultures and resolve very small differences in mutation rate ^60,88^. The *Compare.LD* function was applied pair-wise to all strains, and obtained *p* values were corrected for multiple comparisons by controlling the False Discovery Rate ^89^ using p.adjust in R. Adjusted *p* values less than 0.05 were considered to indicate significant difference in mutation rates.

## 3. Results

Maximum likelihood phylogenetic analysis of 80 *S. iniae* isolates (Table 1) based on 10,267 non-recombinant core genome SNPs derived from whole genome data resolved ten major phylogenetic lineages: six lineages with multiple strains (clades A-F), one lineage with two strains (clade G), and three lineages with a single strain (Figs. 1, 2). Some lineages (C-E) show a degree of geographic endemism while clades (A, B, and F) contain strains from diverse geographic regions. Principally, variation in MMR and GO genes occurs only between the lineages, and they are highly conserved within clades (Fig. 2, Table 2). Most SNPs in MMR and OG genes were non-synonymous and present in protein functional domains, and most amino acid substitutions were predicted to have a deleterious effect on protein function (Table 2). Furthermore, the number of MMR and OG variants correlates significantly with mutation rate (*p* = 0.0056), as predicted by a Phylogenetic Generalised Least Squares (PGLS) regression model that accounts for autocorrelation between the closely related strains ^87^. In turn, mutation rate variation is highly consistent with phylogenetic affiliation: according to multiple pair-wise strain comparisons by Maximum Likelihood Ratio test ^60^, with probabilities corrected for multiple comparison using False Discovery Rate ^89^, in general mutation rate difference is insignificant within but significant between major phylogenetic lineages (Suppl. Table 1). Although mutation rate variation between divergent strains is significant, the magnitude of the differences is modest (around 5-fold) (Figure 1, 2, Suppl. Table 2).

**Table 2:**
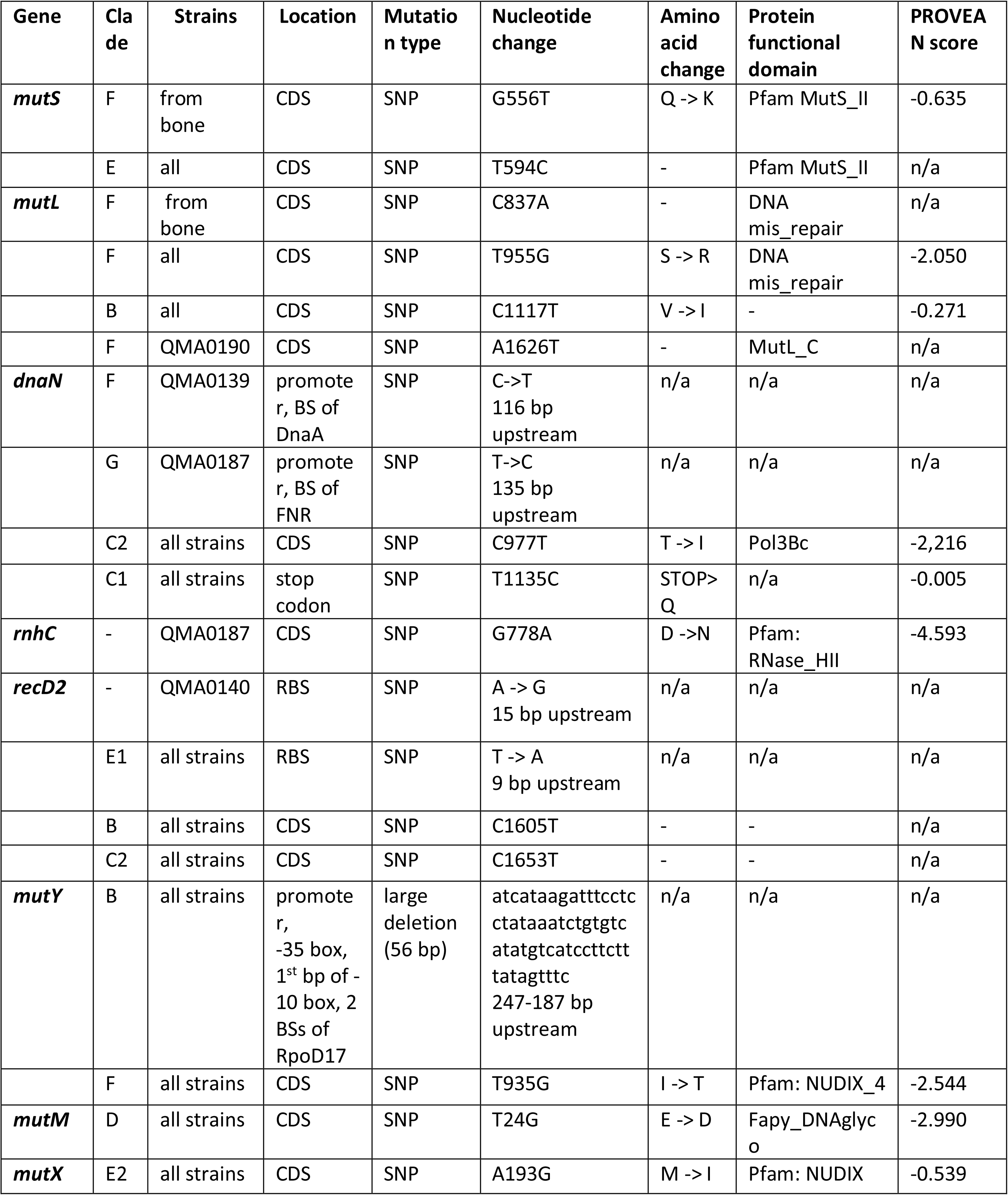
Variants in MMR and OG genes found among *S. iniae* isolates. CDS – protein coding sequence, BS – predicted binding sites of transcription factors, PROVEAN score – predicted effect of amino acid substitutions on the protein function; scores lower that −2 are considered to indicate a deleterious effect.

The accrual of random mutations in MMR/OG loci appears to be the primary mechanism of mutation rate molecular evolution within analysed set of *S. iniae* genomes. Firstly, we used a maximum likelihood inference method to exclude areas of recombination from our phylogenetic analyses ^67^, and none of the MMR and OG gene loci were excluded as recombinant. Within the core genome, where MMR and OG gene loci are located, recombination rates were extremely low with mean μ_r/m_ 0.032 and only 5 from the 80 strains containing recombination blocks (Suppl. Table 5). Secondly, we created a pangenome for the 80 strains in our analysis and mapped on putative phage and prophage signatures ^80^, and insertion sequence elements ^81^. None of the MMR or OG genes occurred in proximity to phage or insertion sequence elements (Suppl. Fig. 2). Also, tandem-repeat copy-number variants identified in *mutL* in *E. coli* ^44,90^ were not detected in MMR or OG genes in *S. iniae* (Table 2). All mutations in MMR/OG loci in our dataset were SNPs, except for the large deletion affecting the *mutY* promoter in clade B (Table 2). As most *S. iniae* lineages contain variants pertaining to both MMR and OG systems (Table 2), the detection of mutational bias produced by either dysfunctional MMR or OG demonstrated in experimental populations of *E.coli* ^91^ is limited for our natural isolates. Nonetheless, consistent with findings in *E. coli* where OG mutants showed increased frequency of AT to CG transversions ^91^, the latter bias was detected in the only *S. iniae* lineage (clade D) where mutations are constrained to a single OG variant (Suppl. Fig. 1). This variant is an amino acid substitution in the DNA glycolyase domain of mutM that is predicted to deleteriously affect the protein function (PROVEAN score of −2.990, Table 2).

To investigate whether variation in mutation rate phenotype and MMR/OG genotype is associated with adaptation to the host, we identified variation in key virulence traits in streptococci (capsular polysaccharide, hemolysin, length of cell chains) and bacteria in general (resistance to ROS, and biofilm formation). We found that number of atypical phenotypes for these traits correlates significantly (*p* = 0.0331, PGLS) with the number of MMR and OG variants (Fig. 2, Table 3). Moreover, infections in atypical hosts (mammals) and tissue within apparently immune primary hosts (bone of vaccinated barramundi) correlates strongly with both number of atypical virulence-associated phenotypes (p < 0.0001) and the number of MMR/OG variants (p = 0.002).

Of the clades identified among *S. iniae*, clade A is apparently a dominant circulating clade primarily infecting Perciform fish that has persisted globally for almost two decades (clade A contains strains isolated from USA, Honduras, and Australia between 1999 to 2016). Mutation rates in all isolates (*n*=35) among this ancestral lineage fall within a 1.5 - 2 × 10^−8^ range (Figs. 1, 2; Clade A), which is similar to the base mutation rate estimated for other non-mutator gram positive bacteria ^11,92^. All differences in mutation rates within this clade are insignificant except three pairwise comparisons that verge on p <0.05 (Suppl. Table 1). MMR and OG genes in these strains are identical, and only minor deviations from wildtype traits contributing to virulence were observed (absence of capsule production and longer cell chains in QMA0158, 216) (Figs. 1- 3, Table 1). All other lineages (with the exceptions of clade C1 and strains QMA0445-46 discussed below) have significantly different mutation rates to clade A, contain unique and often multiple SNPs in MMR/OG loci, and show peculiar phenotypic variants related to virulence (Figs. 1-3, Tables 1, 2, Suppl. Table 1). Although phylogenetic relatedness between lineages is fully resolved (Fig. 2), they originate almost simultaneously, and nested lineages evolve independently sharing little phylogenetic history (Figs. 1, 2). Lack of correlation between branch length and sampling date derived by root-to-tip regression analysis with time is evidence of non-neutral evolution amongst this collection of *S. iniae* isolates (R^2^=0.0936, Correlation coefficient 0.306, best-fit root; Suppl. Fig. 3, ^72^). Deviation from a neutral model is supported by highly skewed branching, with some branches comprising single isolates whilst others comprise many epidemiologically unrelated isolates (Figure 1, 2). Although potentially affected by the sampling bias, this is suggestive of frequent strong selection, presumably imposed by heterogeneity of the immune landscape encountered during transfer between host individuals, and resembles genealogical tree topologies arising from viral evolution over similar timespans ^93,94^.

Clade B contains isolates from ornamental fish, QMA0084 from flying fox, *Epalzeorhynchos kalopterus*, in Western Australia (WA) and QMA0462-63 from clown loach *Chromobotia macracanthus*, in USA. These isolates share a valine to isoleucine substitution in MutL predicted as neutral (PROVEAN score −0.271), a synonymous SNP in *recD2*, and a 56 bp deletion affecting the predicted promoter of *mutY* (Table 2). The latter mutation encompasses the −35 box, two binding sites of the RpoD17 general transcription factor, and the 1^st^ bp of the −10 box, and was predicted to abolish the original promoter (Suppl. Fig. 4). However, *mutY* might still be transcribed at some level since two other potential promoters were identified in a nearby sequence (Suppl. Fig. 3). It appears that altered *mutY* expression is also affected by the rest of the genetic background as isolates exhibit significant differences (Suppl. Table 1) in mutation rate phenotype, 7 × 10^−8^ in QMA0084 and 4 × 10^−8^ in clown loach strains (Figs. 1-2). Phenotypically, clade B has shifted towards decreased haemolytic activity (Fig. 3, B2).

Clade C shares a common ancestor with clade B and is comprised of strains obtained from barramundi farmed in WA, Northern Territory (NT) and north QLD fish farms (nested subclades C1 and C2 respectively). While a core mutation rate of 1.5-2 × 10^−8^ is observed in subclade C1, a significantly higher mutation rate of 7 × 10^−8^ has been maintained in subclade C2 from north QLD for almost two decades (1995 to 2012) (Figs. 1,2 Suppl. Table 1). Minor variation in virulence-related phenotypes (absence of capsule production and longer cell chains in QMA0074, 77) (Fig. 2, Table 1) suggests that evolution in this lineage might be driven by outside-the-host factors. The significant difference in mutation rate between nested clades C1 and C2 may result from variation in *dnaN*. In *Bacillus subtilis*, 90% of mismatch repair is dependent on targeting MutS to nascent DNA via the ß sliding clamp (DnaN) zone ^95,96^. Inactivation of *dnaN1* in *B. anthracis* results in a mutator phenotype with a mutation rate equivalent to a *mutS* mismatch repair-defective strain ^97^. The tyrosine to isoleucine substitution in one of the critical residues of the loader binding interface of the ß-clamp, predicted as deleterious (PROVEAN score −2,216), is present in all isolates of from clade C (Table 2), which may explain the high mutation rate in clade C2 strains (Fig. 2, Table 2). Potentially, rapid fluctuation in water salinity, oxygen and temperature accompanying periodic heavy rainfall in tropical north QLD represent an unstable environment that favours persistence of mutator alleles ^25^. In contrast, only subclade C1 harbours T1135C nucleotide substitution in the *dnaN* that changes stop codon TAA into CAA coding for glutamine predicted as neutral (PROVEAN score −0.005), with the next stop codon TAG found immediately downstream. Putatively, T1135C and/or change of the *dnaN* stop codon ^98^ may act as compensatory to the T326I substitution, reducing the mutation rate to a value not significantly different from the core rate found in clade A (Fig. 1, 2) via protein elongation or by affecting translation via stop codon usage bias ^98^. In addition, a synonymous variant in *recD2* is found in subclade C2 only (Table 2).

Clade D consists of strains from trout (*Oncorhynchus mykiss*) isolated in Réunion and Israel. These strains have an estimated mutation rate of 5 × 10^−8^, contain a glutamate to aspartate substitution in *mutM* predicted to be deleterious to protein function, a synonymous SNP in *RecD2*, and exhibit impeded haemolytic activity (Fig 1,2, Table 2). Also, isolates from Israel form thicker and more dense biofilm structures (Fig. 3, D2).

Isolates from humans and fish are found in clade E where multiple phenotypic profiles for the virulence-associated traits are observed (Figs. 2-3, Table 1). This clade contains two nested subclades that share a SNP in *mutS*, but have other unique MMR/OG SNPs (Fig. 2, Table 2) and exhibit mutation rates that are significantly different in most pairwise comparisons (Suppl. Table 1). Subclade E1 contains USA isolates from humans (QMA0133-35, 37-38) and hybrid striped bass (QMA0447-48), which have a substitution in the Shine-Dalgarno sequence of *recD2* and a mutation rate of 6.5 × 10^−8^ (Fig 1, Table 2). Fish strains and three human strains (QMA0135, 37-38) form shorter cell chains and are less sensitive to H_2_O_2_. Two human strains QMA0133-34 produce thicker and more dense biofilm structures (Fig. 3, D2). QMA0133 is non-encapsulated (Fig 3, A2), and QMA0134 appears to differentially express the capsular polysaccharide in culture (Fig 3, A3). Subclade E2 contains two human strains from Canada (QMA0130-31) and a tilapia strain from USA (QMA0466). These strains have unique methionine to isoleucine substitution at the N-terminus of MutX, and a mutation rate of 4.5 × 10^−8^ (Fig 1, Table 2). Both human strains are non-encapsulated (Fig. 2). In contrast, the tilapia strain expresses the capsule but forms short cell chains in common with most strains from the subclade E1 (Fig 2, Fig 3, C3).

Clade F is not readily definable by location, time of isolation, or host species and comprises QMA0139 from unidentified fish in Canada, QMA0190 isolated from snakehead murrel in Thailand (both of which appear on long branches), and nested terminal clade containing strains QMA0233-36, 249, 254-55 from barramundi farmed in Recirculating Aquaculture Systems (RAS) in New South Wales and South Australia (Fig. 2). The latter were isolated from barramundi bone lesions during a disease outbreak in vaccinated fish where infection manifested itself as a slowly progressing osteomyelitis ^49^ in contrast to typical acute septicaemia and meningitis ^48^. Multiple phenotypic changes associated with chronic infection and increased potential for colonization were present in bone isolates, including absence of capsule, impeded hemolytic activity, increased cell-chain length, and denser biofilms ^53–56,84,99,100^ (Fig. 3). Bone strains have a unique MMR/OG genotype (Fig. 2, Table 2) and a mutation rate that is different from QMA0139 and QMA0190 (Figs. 1, 2, Table 1). A serine to arginine substitution in MutL and tyrosine to isoleucine substitution in MutY seems to have occurred in the ancestor of QMA0139, QMA0190, and bone strains. Both substitutions are predicted to have a deleterious effect on protein function (PROVEAN scores −2.050 and −2.544 respectively), but their effect on mutation rate is combined with unique SNPs found in each branch: QMA0139 has a SNP in the binding site for the DnaA transcription factor within the *dnaN* promoter, and a mutation rate of 4.5 × 10^−8^, QMA0190 has synonymous SNP in *mutL* and a mutation rate of 6.5 × 10^−8^, and bone strains have glutamine to glycine substitution in MutS predicted as neutral (PROVEAN score −0.635), a synonymous SNP in *mutL*, and a mutation rate of 3-4 × 10^−8^. Considering that the mutation rate of QMA0190 is significantly higher compared to rates expressed by QMA0139 and bone isolates (Suppl. Table 1), it is likely that variants unique to each of the latter strains compensate for deleterious variants shared by the isolates ^46^.

Clade G contains two strains from tilapia isolated in USA (QMA0445-46). Notably, despite being phylogenetically distant from clade A, these strains have the same MMR and OG genotype and have retained the core mutation rate and the typical phenotypic profile for virulence-associated traits (with the exception of capsule absence most likely inherited from common ancestor shared with strains from clade F, QMA0140-41, 187 that are also non-encapsulated) (Fig. 2, Table 1).

Three isolates of *S. iniae* were identified as independent phylogenetic lineages, with unique mutation rate genotypes and phenotypes, and virulence traits (Figs. 1-2, Tables 1-2). The first long branch with a single isolate contains a strain from snakehead murrel strain (*Channa striata*) from Thailand. This isolate is non-encapsulated and weakly haemolytic and has a mutation rate of 4.5 × 10^−8^, potentially attributable to a deleterious aspartate to asparagine substitution in rnhC and SNP in the binding site of for the *fnr* transcription factor within the *dnaN* promoter. A second long branch with a single strain contains the oldest among the analysed strains, QMA0140 isolated in 1976 from dolphin (*Inia geoffrensis*) ^101^. This strain exhibits a mutation rate of 1 × 10^−8^, which is unique among the strains and significantly lower than the core mutation rate, perhaps linked to increased translation of recD2 helicase resulting from a SNP in the ribosome binding site (Fig 2, Table 2). The longest branch on the tree contains a second dolphin strain, QMA0141, isolated two years later in 1978 ^102^. This isolate is highly divergent from the rest of strains with around 20 kB of non-recombinant SNPs in pair-wise comparisons with other strains, accounting for around 1% genomic difference. In contrast to QMA140, it mutates at a rate of 1x 10^−7^, the highest mutation rate phenotype determined among the isolates and significantly different to the rest of the values. Multiple SNPs are observed in all MMR and OG genes: 3 in *dnaN* and *mutX*, 4 in *mutM*, 9 in *mutL* and *rnhC*, 15 in *recD2*, 9 in *mutL* and *rnhC*, and 68 in *mutS*. Both dolphin isolates are non-encapsulated, show increased ability to withstand oxidative stress, and form denser and thicker biofilms (Fig 2, 3).

## 4. Discussion

Mutator alleles from MMR and OG systems are abundant among infectious isolates and are likely to promote pathogen evolution since they can greatly increase the supply of pathoadaptive mutations in changing host environments ^11,30,32,37,38^. The dynamics of mutation rate modifiers – increase in mutator alleles in times of adaptation and their decline in adapted communities – is not fully understood and continues to be the focus of multiple experimental studies ^24^. Investigation of such mutation rate dynamics in natural populations is a challenging task considering mutator alleles often revert to wildtype level via back-mutation or recombination/translocation that fully restores the wiltype mutation rate genotype and phenotype ^44,45^. However, when compensatory evolution of mutation rates rather than reversion occurs, i.e. when anti-mutator variants with secondary-site mutations in MMR/OG genes emerge that lower mutation rates – this is likely to produce traceable shifts in DNA repair genotype and phenotype of an adapted strain ^46^. Here we have investigated the mutation rate dynamics in natural populations of the rapidly evolving aquatic pathogen *S. iniae* ^48^, using 80 strains isolates globally over 40 years. We performed whole-genome sequencing, constructed phylogenetic tree based on non-recombinant core genome SNPs ^103^, identified variation in major MMR and OG loci, estimated mutation rates by fluctuation analysis ^58^, and compared the latter by the Likelihood Ratio Test ^51^. As hypothesized, we have found that MMR/OG genotype and mutation rate variation occurs between major phylogenetic lineages and is minimal within phylogenetic clades. While each lineage has unique MMR/OG profile, MMR and OG loci are conserved even within phylogenetic groups not readily definable by geographic origin, time, or host species. Therefore, these loci can be used for rapid diagnostics via Multilocus Sequence Typing (MLST) ^104^. In turn, mutation rates are conserved within phylogenetic clades, while significant differences occur among major phylogenetic lineages. Although significant, differences in mutation rate are largely small (around 5-fold), which is consistent with transience of mutators in natural populations and restoration of low mutation rate after adaptation is gained ^46^. On the other hand, one order of magnitude or lower differences in mutation rate distinguish mutator from non-mutator phenotypes in other streptococcal species ^11,92^.

Arguably, the accrual of local mutations in MMR/OG loci is a primary mechanism of molecular evolution driving mutation rate dynamics in *S. iniae*. Firstly, all variants in MMR and OG genes were single nucleotide substitutions with the exception of the large deletion affecting the *mutY* promoter in clade B. Secondly, we did not find evidence of other mechanisms that underlie the flux of mutator alleles and phenotypes including recombination ^47^, and prophage integration/excision ^45^, and tandem repeat copy-number variation ^44,90^. Indeed, the easily reversible copy-number variation prone to back mutation and full restoration of the wiltype non-mutator genotype and phenotype, to date has been only found in *mutL* in *Proteobacteria* ^44,90^, which is structurally and functionally different from its homologs in gram-positive bacteria and eukaryotes ^19,52^. Notably, a single SNP in MMR or OG loci can be sufficient to induce profound changes in mutation rate ^105,106^, and most of the SNPs identified in our isolates were found within coding regions, non-synonymous, and predicted to negatively affect the protein function. The dysfunctional MMR and OG can leave molecular signatures within the genome in the form of mutational bias: as observed in a long term evolutionary experiment in *E.coli* defective MMR and OG repair increased frequency of AT to GC transitions and AT to CG transversions respectively ^91^. Such analysis is limited for our dataset since most of our natural *S. iniae* isolates contain both MMR and OG variants. Nonetheless, we quantified the mutational spectra by clade, and, consistent with findings in *E. coli* experimental mutators, the highest frequency of AT to CG transversions was observed in the only lineage containing a single OG variant (clade D with the amino acid substitution in MutM in predicted as deleterious). Thus, it appears that mutation rate dynamics in *S. iniae is* primarily driven by local mutations and compensatory mutator/anti-mutator evolution where some variants increase and some decrease the mutation rate ^46^.

The conservation of MMR/OG genotypes within *S. iniae* clades and their variation among the major phylogenetic lineages, linkage of unique MMR/OG profiles to unique mutation rate values, and the likelihood of most identified SNPs to affect protein function or translation rate are consistent with an idea that mutator phenotypes may be facilitating adaptations preceding major diversification events in *S. iniae* evolution. The tree topology with highly skewed branching deviates from a neutral model and resembles a genealogical tree topology derived from viral evolution over similar timespans, suggesting evolution under strong selection ^93,94^. Since selection by the host immune responses is the major force shaping strain diversity in pathogens ^6^, we phenotyped our strains for major traits determining virulence in streptococci - capsular polysaccharide, hemolysin, length of cell chains, resistance to reactive oxygen species, and biofilm formation^53–57^. As estimated by a Phylogenetic Least Squares (PGLS) model accounting for autocorrelation among phylogenetically close strains ^87^, the occurrence of non-typical phenotypes for these traits correlates with number of MMR/OG variants. The variation in virulence phenotypes is minimal and relatively low mutation rate (core rate of 1.5-2 × 10^−8^) is observed in apparently dominant globally distributed clade A and also clade G with MMR/OG profile and mutation rate identical to clade A. Other lineages contain MMR/OG variants, express 3-5 times higher mutation rates (except mutator strain from dolphin with 10 times higher mutation rate), and contain isolates with non-typical virulence-associated variants. This is consistent with an idea of selection for lower mutation rates in adapted populations ^46^. One the other hand, in contrast to other common pathogens where 2-3 orders of magnitude differences in mutation rate are observed between mutators and non-mutators ^11^, only one order of magnitude or lower differences appear to distinguish mutator from non-mutators in streptococci (around 10^−7^ and 10^−8^ mutation rates respectively) ^11,92^.

Furthermore, the most variants in MMR/OG SNPs and changes in virulence phenome are observed in strains from atypical immune settings - atypical hosts (mammals) and atypical tissue within apparently immune vaccinated fish hosts (barramundi bone). Firstly, this implies that mutators may promote pathogen transmission to novel hosts (’host jumps’), including transmission between the higher host taxa such as from fish to mammals discussed in the present study ^107^. The adaptation to an ultimately different immune responses of a novel host species is likely to require multiple genetic and phenotypic variants, whose occurrence on the same genetic background is most readily achievable by the mutator strains. Therefore, it is conceivable that transfers from wild fish to dolphins (strains QMA0140-41) and farmed hybrid bass to humans ^108^ in lineage E have been facilitated by the mutator phenotype. Second, mutators might present a risk factor to immune escape after vaccination and vaccine-induced serotype replacement (VIRS) – the spread of co-existing pathogen serotypes after elimination of the dominant serotypes vaccinated against ^109^. An atypical pathology outbreak caused by strains from clade F has occurred on RAS in New South Wales and South Australia where fish were immunised using whole-cell killed bacterins prepared from strains found in dominant clade A ^49^. Notably, vaccine-induced immunity was partially effective in the infected fish, evident by cross-reactivity of vaccine-induced antibodies against clade F strains and absence of their proliferation in typical niduses (blood, brain, and pronephros) ^49^. Nonetheless, colonization of the barramundi bone tissue by clade F strains caused slowly progressing osteomyelitis instead of typical acute septicaemia and meningitis ^48,49^. These bone isolates express multiple non-typical phenotypes - absence of capsule, impeded hemolytic activity, increased cell-chain length, and denser biofilms. All these shifts are likely to be adaptive as they were previously associated with chronic infection and increased potential for colonization ^53–56,84,99,100^. Multiplicity of virulence-associate phenotypic shifts in bone strains indicates that evasion from the immune clearance and adaptation to osseous tissue are likely to have occurred via the mutator phenotype. Indeed, bone isolates have unique mutation rates and multiple MMR and OG SNPs where some variants appear to have mutator and some anti-mutator effect. Thus, mutator alleles may potentially promote evolution of novel serotypes and VISR in populations of farmed fish. Notably, as evidenced by bone isolates, VIRS might occur without major disruption of immune recognition when multiple changes permit colonisation of and persistence in the tissue with lower immune surveillance. Extension to other species such as *Streptococcus pneumoniae* may be particularly interesting as serotype evolution following vaccination programmes is well documented, but the role of mutators in this process is unknown ^110–112^. Since mutators may promote emergence of novel serotypes, development of long-term efficient generic polyvalent vaccine protective against all co-existing strains may not be readily achievable with rapidly evolving microorganisms. Thus, a flexible approach to veterinary vaccine licencing and re-formulation may be recommended to locally eliminate the emergent virulent serotypes of antigenically variable pathogens before they spread in the population.

In summary, our results broadly support the notion that mutators promote evolution of virulence and antigenicity, leading to major diversification events in *S. iniae*. More specifically, they may facilitate adaptation to a new immunological setting, such as one required for exploitation of a new host taxon, new host tissue/s, or a host with vaccine-induced adaptive immune response. Therefore, dynamics of mutator alleles in natural populations of pathogens could be associated with epidemiologically relevant incidents such as vaccine escape outbreaks and host jumps that should be explored further.

## Acknowledgements

This work was supported by The Australian Research Council Discovery Project Grant DP120102755 awarded to ACB. JE was supported by an ARC Future Fellowship (FT140100907). We thank the University of Queensland School of Biological Sciences for funding a tuition fee waiver for OS. For provision of strains into our collection over 15 years we thank: Judy Forbes-Faulkner, Oonoonba Veterinary Laboratory, QLD, Australia; Lynn Shewmaker, CDC, Atlanta, USA; Christian Michel, INRA, France; Suresh Benedict, Berrimah Veterinary Laboratories, NT, Australia; Nicky Buller, DPIRD WA, WA, Australia; Mark White, Tréidlia Biovet Pty (formerly Allied Biotechnology), NSW, Australia; Matt Landos, Future Fisheries Veterinary Services, NSW, Australia; Spencer Russell, Novartis Animal Health, Canada; Ruben Arturo Lopez Crespo, Regal Springs Tilapia, Honduras. We thank Dr Simone Blomberg for advice on statistical analyses and Mr Nazar Rudenko for assistance with writing software pipelines and for use of his Unix server.

---

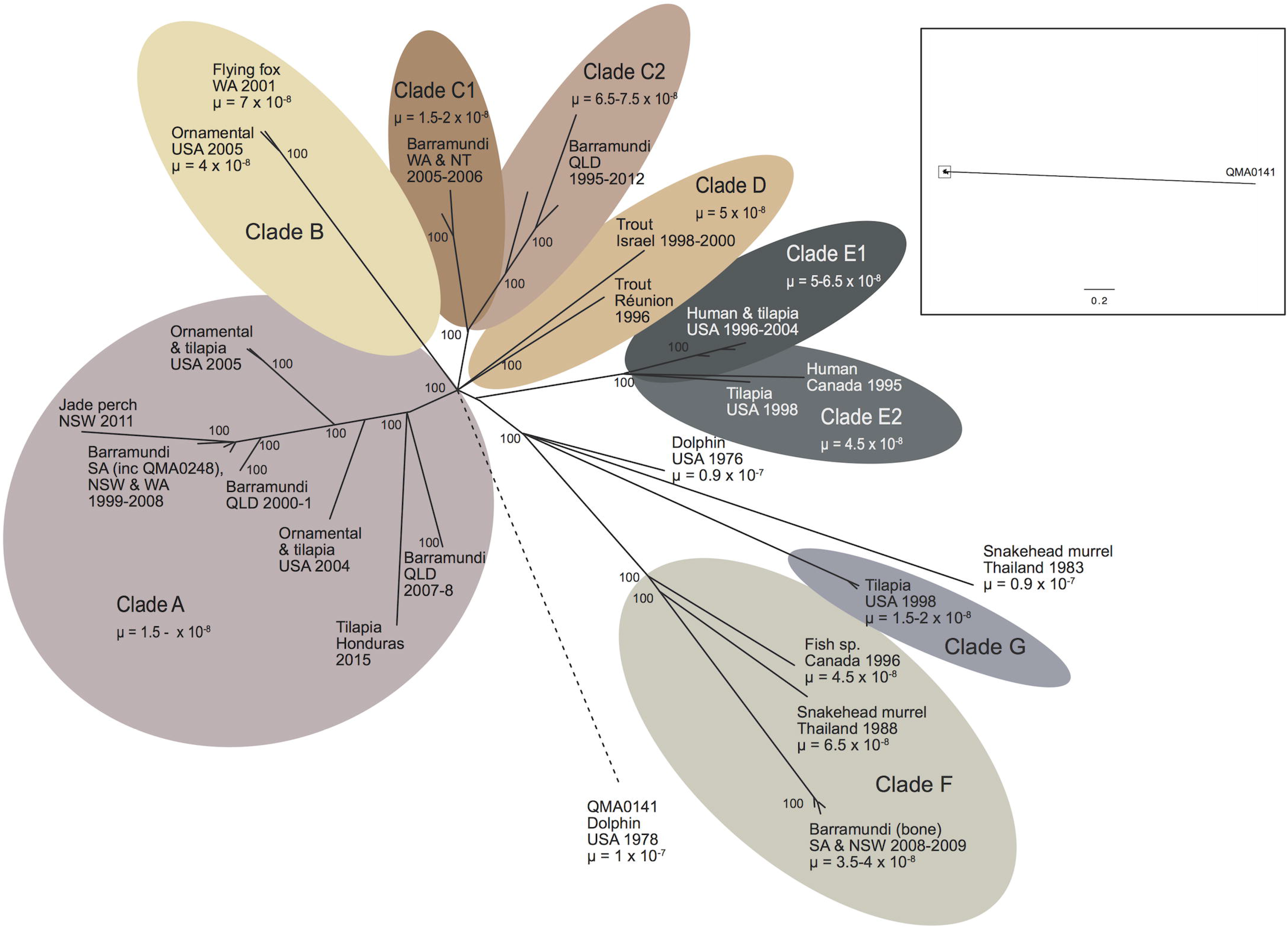

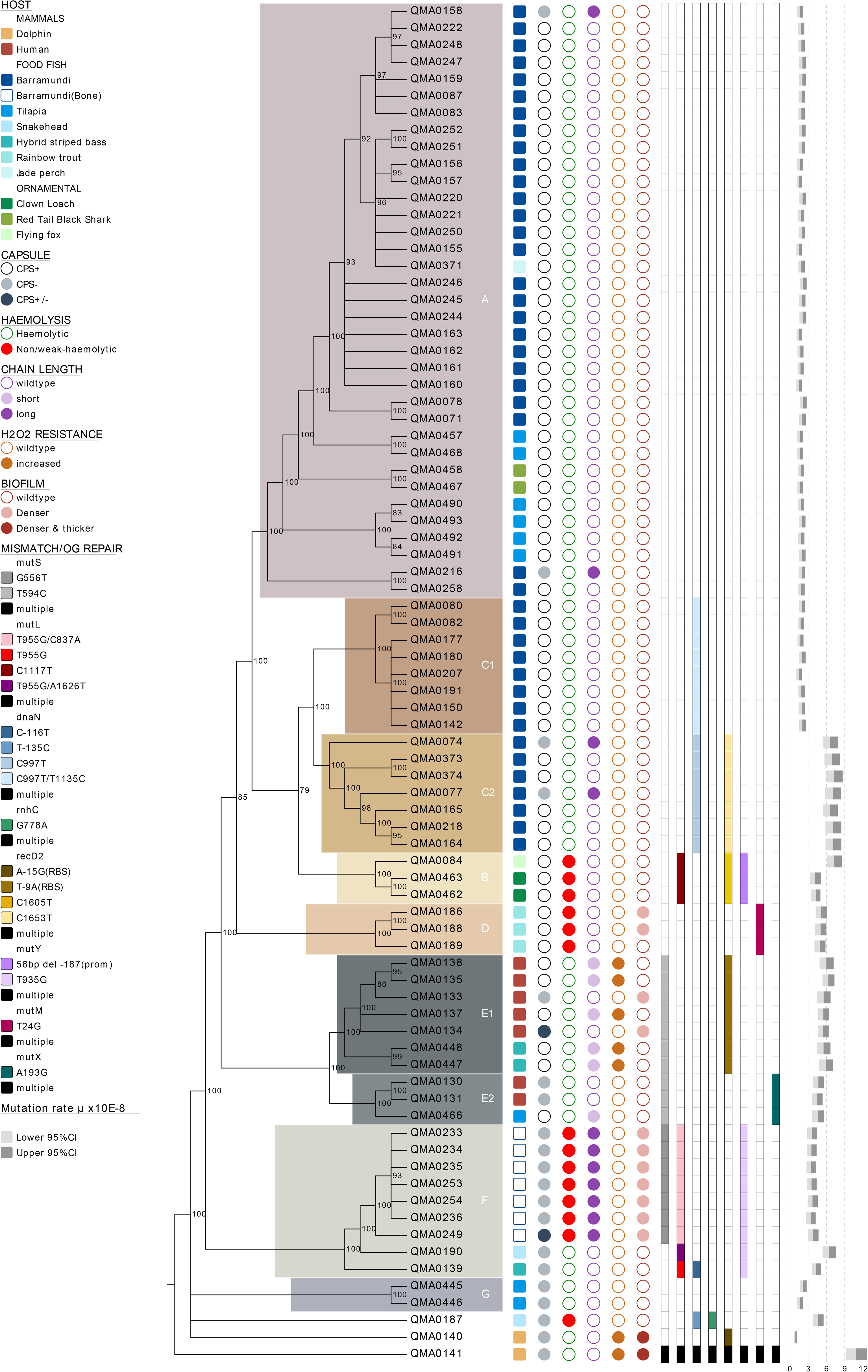

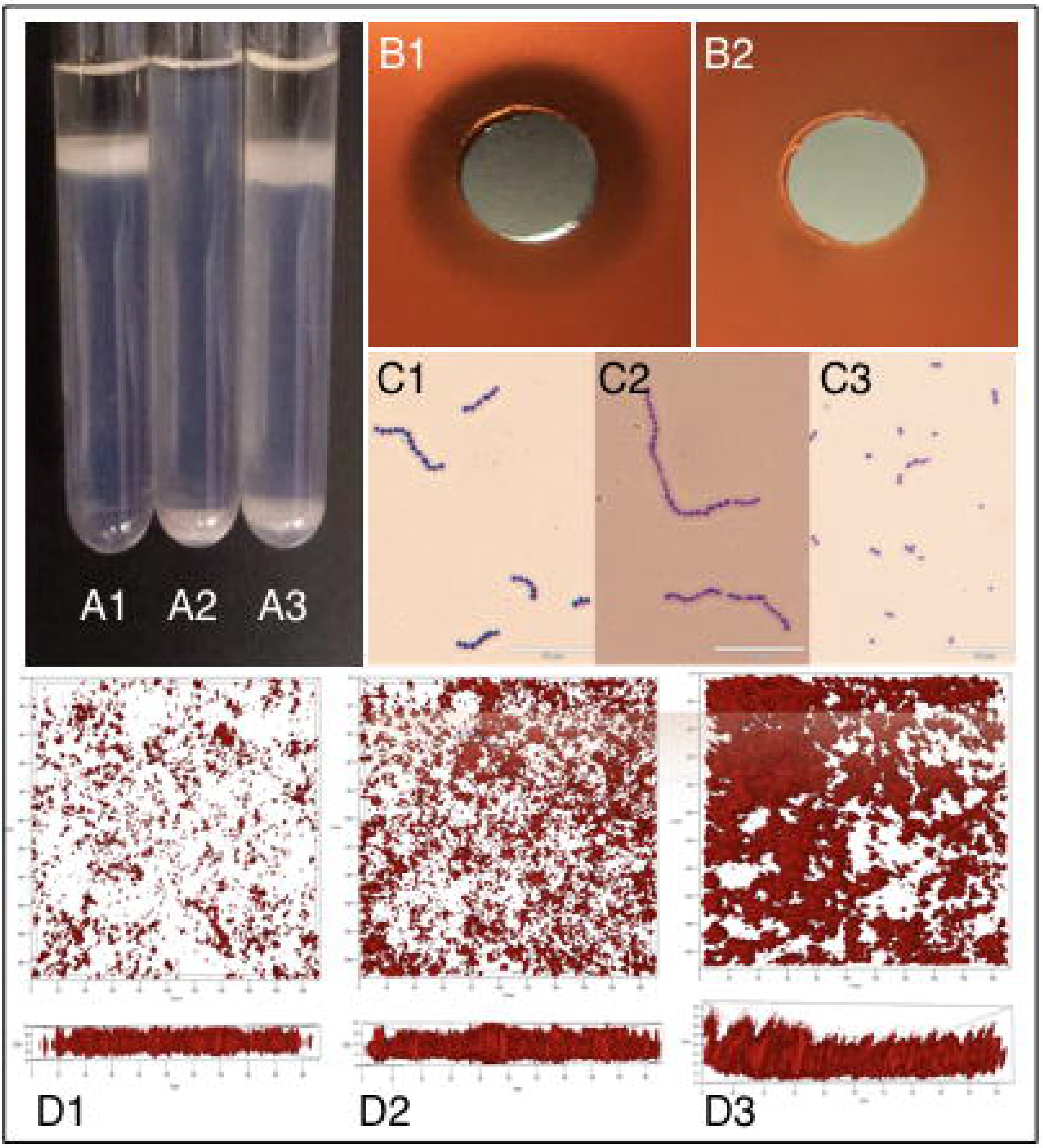

