## Supplementary figures and images for "Evolutionary epidemiology of *Streptococcus iniae*: linking mutation rate dynamics with adaptation to novel immunological landscapes"

### Supplementary Fig 1

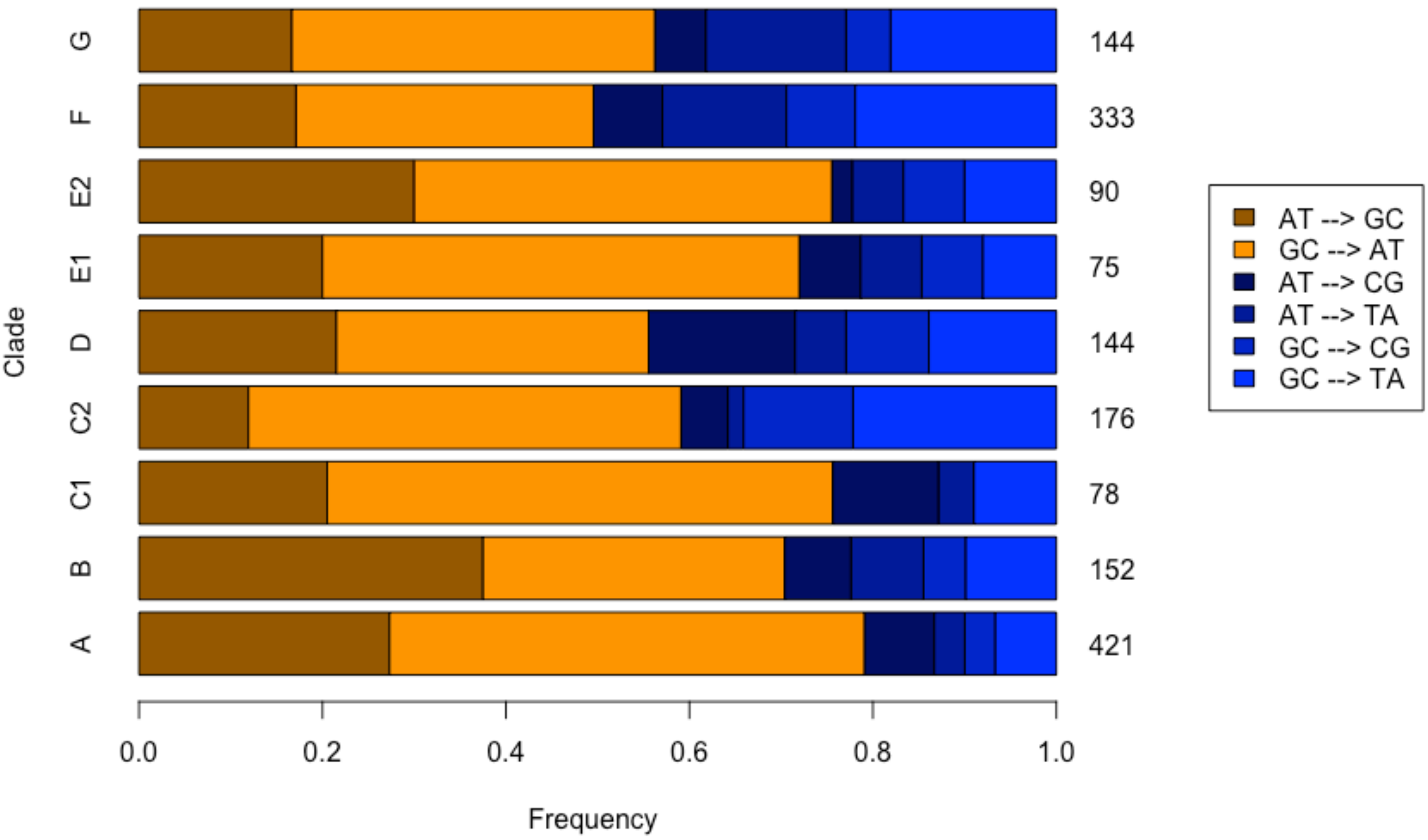

### Supplementary Fig 2

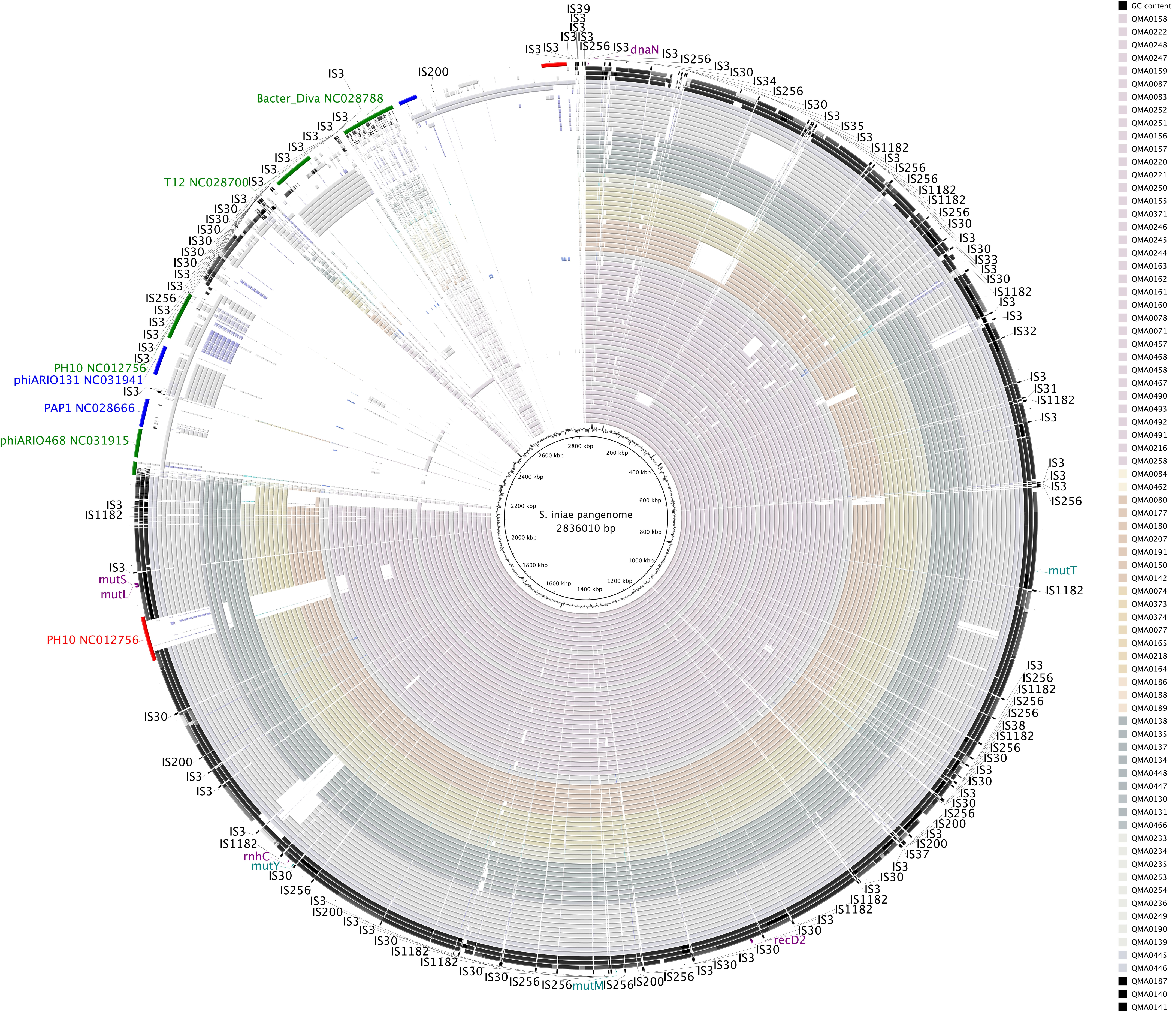
