## Supplementary Fig 3 for "Evolutionary epidemiology of *Streptococcus iniae*: linking mutation rate dynamics with adaptation to novel immunological landscapes"

☑ Best-fitting root

Function: heuristic resid...

| Dated Tips | |
|---|---|
| Date range | 39 |
| Slope (rate) | 4.7079E-4 |
| X-Intercept (T... | 1822.6921 |
| Correlation Co... | 0.3059 |
| R squared | 9.36E-2 |
| Residual Mean... | 9.8948E-5 |

Sample Dates | Tree | Root-to-tip | Residuals | Node density

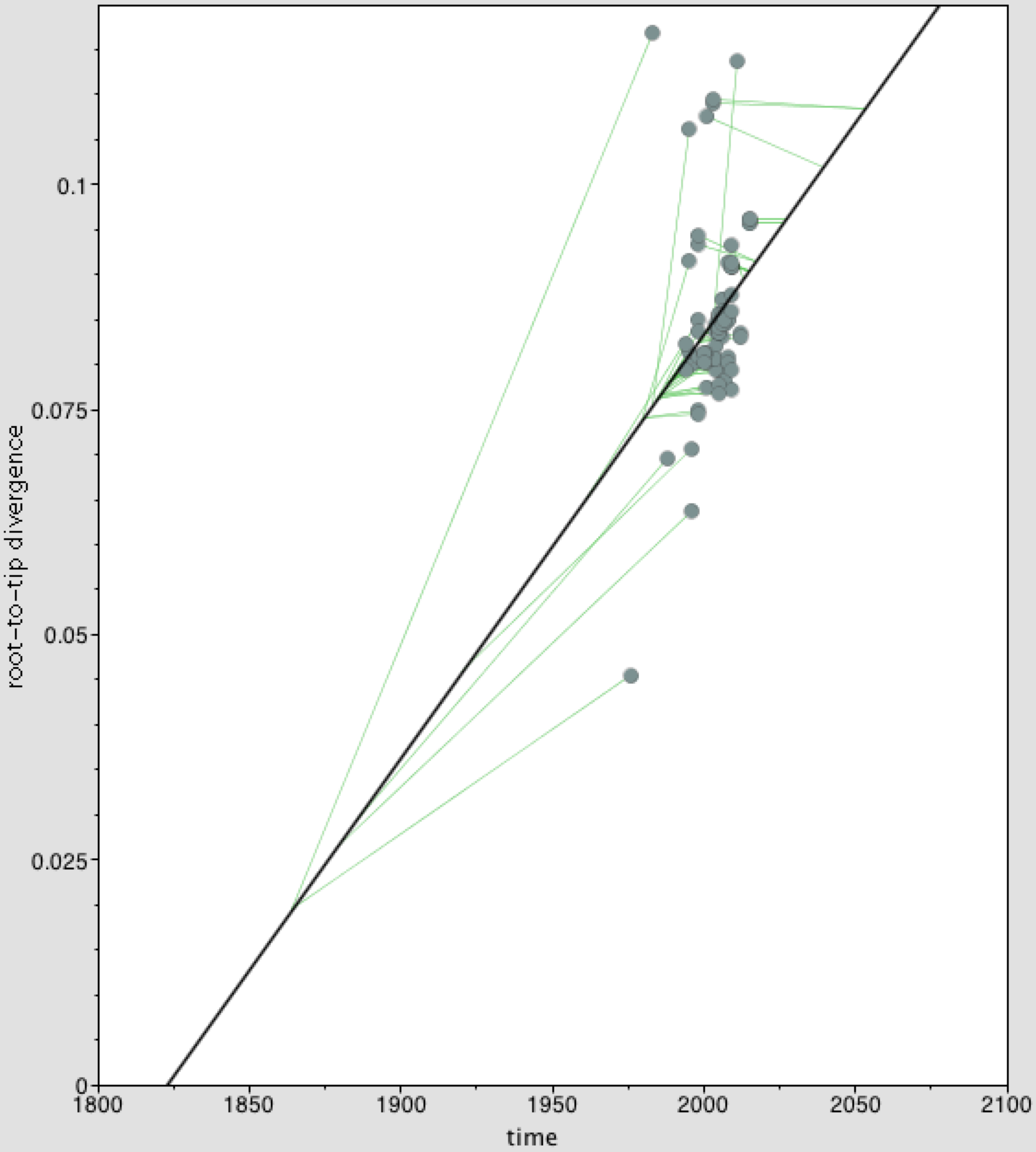
