## Supplementary Fig 4 for "Evolutionary epidemiology of *Streptococcus iniae*: linking mutation rate dynamics with adaptation to novel immunological landscapes"

```
QMA248_500      cagattatgaataactaatctctggaaaaattagtctccattcttgtaacgtcctttatt   60
QMA0084_444     cagattatgaataactaatctctggaaaaattagtctccattcttgtaacgtcctttatt   60
                ************************************************************

QMA248_500      gaatttaagcatatactatacctttcaatattggagattttccaaaacttatactgtaaa   120
QMA0084_444     gaatttaagcatatactatacctttcaatattggagattttccaaaacttatactgtaaa   120
                ************************************************************

QMA248_500      gagcgctcctttggccaaataacgcatacttcaataagttcaacaatgtgatcactttgt   180
QMA0084_444     gagcgctcctttggccaaataacgcatacttcaataagttcaacaatgtgatcactttgt   180
                ************************************************************

QMA248_500      tagttgtaaaagtcagtttgcctaccgttgttggagctacttagtctccgttcttgtacc   240
QMA0084_444     tagttgtaaaagtcagtttgcctaccgttgttggagctacttagtctccgttcttgtacc   240
                ************************************************************

QMA248_500      gtgcgacatgtagtttcatcataagatttcctcctataaatctgtgtcatatgtcatcct   300
QMA0084_444     gtgcgacatgtagtttc-------------------------------------------   257
                *****************

QMA248_500      tctttatagtttcattataatgtaagcgtttgcttttgtcaagaaaaaatatttccctat   360
QMA0084_444     -------------attataatgtaagcgtttgcttttgtcaagaaaaaatatttccctat   304
                             ***********************************************

QMA248_500      gacattttccacttgcctaataaaaagataagaaaaaaaagctgttatcattgtttttg   420
QMA0084_444     gacattttccacttgcctaataaaaagataagaaaaaaaagctgttatcattgtttttg   364
                ************************************************************

QMA248_500      aaaactctgatagaagctattattttgactttctagcctagccttttcgcatattgaaaa   480
QMA0084_444     aaaactctgatagaagctattattttgactttctagcctagccttttcgcatattgaaaa   424
                ************************************************************

QMA248_500      aactttgctacaatagaggt   500
QMA0084_444     aactttgctacaatagaggt   444
                ********************
```

**BROM prediction:**

***> QMA248_500***

Length of sequence- 500
Threshold for promoters - 0.20

Number of predicted promoters - 1

Promoter Pos: 328 LDF- 7.23
10 box at pos. 313 cattataat Score 75
35 box at pos. 291 atgtca Score 23

Oligonucleotides from known TF binding sites:

For promoter at 328:
rpoD17: TTCCTCCT at position 268 Score - 8
rpoD17: GTGTCATA at position 284 Score - 7
rpoD17: TAATGTAA at position 318 Score - 12
galR: TGTAAGCG at position 321 Score - 8
rpoD18: TTGCTTTT at position 330 Score - 6
rpoD16: TGTCAAGA at position 337 Score - 19

***> QMA0084_444***

Length of sequence- 444
Threshold for promoters - 0.20

Number of predicted promoters - 2

Promoter Pos: 362 LDF- 6.79
10 box at pos. 347 tgttatcat Score 75
35 box at pos. 324 ataaaa Score 7

Promoter Pos: 62 LDF- 3.55
10 box at pos. 44 ttgtaacgt Score 38
35 box at pos. 23 tggaaa Score 23

Oligonucleotides from known TF binding sites:

For promoter at 362:
arcA: AATAAAAA at position 323 Score - 12
purR: ATAAAAAG at position 324 Score - 11
cpxR: TAAAAAGA at position 325 Score - 9
rpoD18: ATCATTGT at position 351 Score - 5
rpoD19: ATTGTTTT at position 354 Score - 7
fnr: TTTTTTGA at position 358 Score - 9

For promoter at 62:
lrp: ATAACTAA at position 11 Score - 14
tus: TAACTAAT at position 12 Score - 17
tus: TTGTAACG at position 44 Score - 7
rpoD17: ATACTATA at position 73 Score - 11
