## Supplementary Table 3 for "Evolutionary epidemiology of *Streptococcus iniae*: linking mutation rate dynamics with adaptation to novel immunological landscapes"

| Collection na | Other designatio | Biosample Number | Species | Host | Isolation date | Geographic location | Received from | Case notes | References |
|---|---|---|---|---|---|---|---|---|---|
| QMA0071 | *0-43154* | SAMN09104459 | *Streptococcus iniae* | *Lates calcarifer* | 2000 | Australia: QLD | Oonoonba Veterinary Laboratory | FW farm | |
| QMA0074 | *98-51343* | SAMN09104460 | *Streptococcus iniae* | *Lates calcarifer* | 1998 | Australia: QLD | Oonoonba Veterinary Laboratory | FW farm | R. A. Nawawi, J. C. F. Baiano, A. C. Barnes, *Journal of Fish Diseases* **31**, 305-309 (2008). |
| QMA0077 | *95-47554* | SAMN09104461 | *Streptococcus iniae* | *Lates calcarifer* | 1995 | Australia: QLD | Oonoonba Veterinary Laboratory | Marine farm | |
| QMA0078 | *1-41291* | SAMN09104462 | *Streptococcus iniae* | *Lates calcarifer* | 2001 | Australia: QLD | Oonoonba Veterinary Laboratory | FW farm | |
| QMA0080 | *AS-04-1503#7* | SAMN09104463 | *Streptococcus iniae* | *Lates calcarifer* | 2004 | Australia: WA | WA Fisheries | FW farm | R. A. Nawawi, J. C. F. Baiano, A. C. Barnes, *Journal of Fish Diseases* **31**, 305-309 (2008). |
| QMA0082 | *AS-04-0018#1* | SAMN09104464 | *Streptococcus iniae* | *Lates calcarifer* | 2004 | Australia: WA | WA Fisheries | FW farm | |
| QMA0083 | *AS-04-0006#1* | SAMN09104465 | *Streptococcus iniae* | *Lates calcarifer* | 2004 | Australia: WA | WA Fisheries | FW farm | C. M. Millard *et al.* , *Appl Environ Microbiol* **78**, 8219-8226 (2012). |
| QMA0084 | *AS-01-2199#1* | SAMN09104466 | *Streptococcus iniae* | *Pteropus alecto* | 2001 | Australia: WA | WA Fisheries | Black flying fox | R. A. Nawawi, J. C. F. Baiano, A. C. Barnes, *Journal of Fish Diseases* **31**, 305-309 (2008). |
| QMA0087 | *Lake Argyll 1* | SAMN09104467 | *Streptococcus iniae* | *Lates calcarifer* | 2004 | Australia: WA | WA Fisheries | FW farm | R. A. Nawawi, J. C. F. Baiano, A. C. Barnes, Journal of Fish Diseases 31, 305-309 (2008). |
| QMA0130 | *ss1440* | SAMN09104468 | *Streptococcus iniae* | *Homo sapiens* | 1995 | Canada | CDC, Atlanta | | R. Facklam, J. Elliott, L. Shewmaker, A. Reingold, *Journal of Clinical Microbiology* **43**, 933-937 (2005). |
| QMA0131 | *ss1543* | SAMN09104469 | *Streptococcus iniae* | *Homo sapiens* | 1995 | Canada | CDC, Atlanta | | R. Facklam, J. Elliott, L. Shewmaker, A. Reingold, Journal of Clinical Microbiology 43, 933-937 (2005). |
| QMA0133 | *4780-01* | SAMN09104470 | *Streptococcus iniae* | *Homo sapiens* | 2001 | USA | CDC, Atlanta | | R. Facklam, J. Elliott, L. Shewmaker, A. Reingold, Journal of Clinical Microbiology 43, 933-937 (2005). |
| QMA0134 | *4787-01* | SAMN09104471 | *Streptococcus iniae* | *Homo sapiens* | 2001 | USA | CDC, Atlanta | | R. Facklam, J. Elliott, L. Shewmaker, A. Reingold, Journal of Clinical Microbiology 43, 933-937 (2005). |
| QMA0135 | *2388-02* | SAMN09104472 | *Streptococcus iniae* | *Homo sapiens* | 2002 | USA | CDC, Atlanta | | R. Facklam, J. Elliott, L. Shewmaker, A. Reingold, Journal of Clinical Microbiology 43, 933-937 (2005). |
| QMA0137 | *105-04* | SAMN09104473 | *Streptococcus iniae* | *Homo sapiens* | 2004 | USA | CDC, Atlanta | | R. Facklam, J. Elliott, L. Shewmaker, A. Reingold, Journal of Clinical Microbiology 43, 933-937 (2005). |
| QMA0138 | *4989-04* | SAMN09104474 | *Streptococcus iniae* | *Homo sapiens* | 2004 | USA | CDC, Atlanta | | R. Facklam, J. Elliott, L. Shewmaker, A. Reingold, Journal of Clinical Microbiology 43, 933-937 (2005). |
| QMA0139 | *2880-96* | SAMN09104475 | *Streptococcus iniae* | *Fish sp.* | 1996 | Canada | CDC, Atlanta | | R. Facklam, J. Elliott, L. Shewmaker, A. Reingold, Journal of Clinical Microbiology 43, 933-937 (2005). |
| QMA0140 | *ss1056* | SAMN09104476 | *Streptococcus iniae* | *Inia geoffrensis* | 1976 | USA | CDC, Atlanta | | R. Facklam, J. Elliott, L. Shewmaker, A. Reingold, Journal of Clinical Microbiology 43, 933-937 (2005). |
| QMA0141 | *ss1123* | SAMN09104477 | *Streptococcus iniae* | *Inia geoffrensis* | 1978 | USA | CDC, Atlanta | | R. Facklam, J. Elliott, L. Shewmaker, A. Reingold, Journal of Clinical Microbiology 43, 933-937 (2005). |
| QMA0142 | 05/0430-D | SAMN09104478 | *Streptococcus iniae* | *Lates calcarifer* | 2005 | Australia: NT | Berrimah Veterinary Laboratories | isolated from non- vaccinated fish | C. M. Millard et al., Appl Environ Microbiol 78, 8219-8226 (2012). |
| QMA0150 | 05/1151-D | SAMN09104479 | *Streptococcus iniae* | *Lates calcarifer* | 2005 | Australia: NT | Berrimah Veterinary Laboratories | isolated from non- vaccinated fish | C. M. Millard et al., Appl Environ Microbiol 78, 8219-8226 (2012). |
| QMA0155 | 1 | SAMN09104480 | *Streptococcus iniae* | *Lates calcarifer* | 2005 | Australia: NSW | Allied Biotechnology Pty LTd | Isolated from diseased fish in RAS | C. M. Millard et al., Appl Environ Microbiol 78, 8219-8226 (2012). |
| QMA0156 | 2 | SAMN09104481 | *Streptococcus iniae* | *Lates calcarifer* | 2005 | Australia: NSW | Allied Biotechnology Pty LTd | Isolated from diseased fish in RAS | R. A. Nawawi, J. C. F. Baiano, A. C. Barnes, Journal of Fish Diseases 31, 305-309 (2008). |
| QMA0157 | 3 | SAMN09104482 | *Streptococcus iniae* | *Lates calcarifer* | 2005 | Australia: NSW | Allied Biotechnology Pty LTd | Isolated from diseased fish in RAS | R. A. Nawawi, J. C. F. Baiano, A. C. Barnes, Journal of Fish Diseases 31, 305-309 (2008). |
| QMA0158 | 4 | SAMN09104483 | *Streptococcus iniae* | *Lates calcarifer* | 2006 | Australia: SA | Allied Biotechnology Pty LTd | Isolated from diseased fish in RAS | R. A. Nawawi, J. C. F. Baiano, A. C. Barnes, Journal of Fish Diseases 31, 305-309 (2008). |
| QMA0159 | 5 | SAMN09104484 | *Streptococcus iniae* | *Lates calcarifer* | 2006 | Australia: SA | Allied Biotechnology Pty LTd | Isolated from diseased fish in RAS | C. M. Millard et al., Appl Environ Microbiol 78, 8219-8226 (2012). |
| QMA0160 | 6 | SAMN09104485 | *Streptococcus iniae* | *Lates calcarifer* | 1999 | Australia: SA | Allied Biotechnology Pty LTd | Isolated from diseased fish in RAS | C. M. Millard et al., Appl Environ Microbiol 78, 8219-8226 (2012). |
| QMA0161 | | SAMN09104486 | *Streptococcus iniae* | *Lates calcarifer* | 2000 | Australia: SA | Allied Biotechnology Pty LTd | Isolated from diseased fish in RAS | |
| QMA0162 | | SAMN09104487 | *Streptococcus iniae* | *Lates calcarifer* | 2000 | Australia: SA | Allied Biotechnology Pty LTd | Isolated from diseased fish in RAS | |
| QMA0163 | | SAMN09104488 | *Streptococcus iniae* | *Lates calcarifer* | 2000 | Australia: SA | Allied Biotechnology Pty LTd | Isolated from diseased fish in RAS | |
| QMA0164 | 06-40191 | SAMN09104489 | *Streptococcus iniae* | *Lates calcarifer* | 2006 | Australia: QLD | Oonoonba Veterinary Laboratory | | R. A. Nawawi, J. C. F. Baiano, A. C. Barnes, Journal of Fish Diseases 31, 305-309 (2008). |
| QMA0165 | 06-40593 | SAMN09104490 | *Streptococcus iniae* | *Lates calcarifer* | 2006 | Australia: QLD | Oonoonba Veterinary Laboratory | | C. M. Millard et al., Appl Environ Microbiol 78, 8219-8226 (2012). |
| QMA0177 | 06/0784-G | SAMN09104491 | *Streptococcus iniae* | *Lates calcarifer* | 2006 | Australia: NT | Berrimah Veterinary Laboratories | isolated from vaccinated fish against NT1 | C. M. Millard et al., Appl Environ Microbiol 78, 8219-8226 (2012). |
| QMA0180 | 06/0784-M | SAMN09104492 | *Streptococcus iniae* | *Lates calcarifer* | 2006 | Australia: NT | Berrimah Veterinary Laboratories | isolated from vaccinated fish against NT1 | |
| QMA0186 | KFP404 | SAMN09104493 | *Streptococcus iniae* | *Oncorhynchus mykiss* | 2000 | Israel | Dr Christian Michel, INRA, France | | C. M. Millard et al., Appl Environ Microbiol 78, 8219-8226 (2012). |
| QMA0187 | 108-83 (4) | SAMN09104494 | *Streptococcus iniae* | *Channa striata* | 1983 | Thailand | Dr Christian Michel, INRA, France | | R. A. Nawawi, J. C. F. Baiano, A. C. Barnes, Journal of Fish Diseases 31, 305-309 (2008). |
| QMA0188 | KFP173 | SAMN09104495 | *Streptococcus iniae* | *Oncorhynchus mykiss* | 1998 | Israel | Dr Christian Michel, INRA, France | | C. M. Millard et al., Appl Environ Microbiol 78, 8219-8226 (2012). |
| QMA0189 | 21-96 (2) | SAMN09104496 | *Streptococcus iniae* | *Oncorhynchus mykiss* | 1996 | Reunion | Dr Christian Michel, INRA, France | | R. A. Nawawi, J. C. F. Baiano, A. C. Barnes, Journal of Fish Diseases 31, 305-309 (2008). |
| QMA0190 | CII.5b-88 | SAMN09104497 | *Streptococcus iniae* | *Channa striata* | 1988 | Thailand | Dr Christian Michel, INRA, France | | R. A. Nawawi, J. C. F. Baiano, A. C. Barnes, Journal of Fish Diseases 31, 305-309 (2008). |
| QMA0191 | *05/0409* | SAMN09104498 | *Streptococcus iniae* | *Lates calcarifer* | 2005 | Australia: NT | Berrimah Veterinary Laboratories | Strain in autogenous vaccine | C. M. Millard et al., Appl Environ Microbiol 78, 8219-8226 (2012). |
| QMA0207 | *06/0064-I* | SAMN09104499 | *Streptococcus iniae* | *Lates calcarifer* | 2006 | Australia: NT | Berrimah Veterinary Laboratories | Isolated from vaccinated fish | |
| QMA0216 | E | SAMN09104500 | *Streptococcus iniae* | *Lates calcarifer* | 2007 | Australia: QLD | Allied Biotechnology Pty LTd | Isolated from diseased fish in earthen ponds | |
| QMA0218 | G | SAMN09104501 | *Streptococcus iniae* | *Lates calcarifer* | 2007 | Australia: QLD | Allied Biotechnology Pty LTd | Isolated from diseased fish in earthen ponds | C. M. Millard et al., Appl Environ Microbiol 78, 8219-8226 (2012). |
| QMA0220 | I | SAMN09104502 | *Streptococcus iniae* | *Lates calcarifer* | 2006 | Australia: NSW | Allied Biotechnology Pty LTd | Isolated from diseased fish in RAS | |
| QMA0221 | J | SAMN09104503 | *Streptococcus iniae* | *Lates calcarifer* | 2007 | Australia: NSW | Allied Biotechnology Pty LTd | Isolated from diseased fish in RAS | |
| QMA0222 | K | SAMN09104504 | *Streptococcus iniae* | *Lates calcarifer* | 2006 | Australia: SA | Allied Biotechnology Pty LTd | Isolated from diseased fish in RAS | |
| QMA0233 | 1 | SAMN09104505 | *Streptococcus iniae* | *Lates calcarifer* | 2008 | Australia: NSW | Allied Biotechnology Pty LTd | Nov. batch (Harvest, fistulous Pract, spinal osteo | C. M. Millard et al., Appl Environ Microbiol 78, 8219-8226 (2012). |
| QMA0234 | 2 | SAMN09104506 | *Streptococcus iniae* | *Lates calcarifer* | 2009 | Australia: NSW | Allied Biotechnology Pty LTd | bone isolate, March batch | |
| QMA0235 | 3 | SAMN09104507 | *Streptococcus iniae* | *Lates calcarifer* | 2009 | Australia: NSW | Allied Biotechnology Pty LTd | bone isolate, March batch | |
| QMA0236 | 4 | SAMN09104508 | *Streptococcus iniae* | *Lates calcarifer* | 2009 | Australia: NSW | Allied Biotechnology Pty LTd | bone isolate, March 09 | C. M. Millard et al., Appl Environ Microbiol 78, 8219-8226 (2012). |
| QMA0244 | R2 | SAMN09104509 | *Streptococcus iniae* | *Lates calcarifer* | 2008 | Australia: SA | Allied Biotechnology Pty LTd | Isolated from diseased fish in RAS | C. M. Millard et al., Appl Environ Microbiol 78, 8219-8226 (2012). |
| QMA0245 | R3 | SAMN09104510 | *Streptococcus iniae* | *Lates calcarifer* | 2008 | Australia: SA | Allied Biotechnology Pty LTd | Isolated from diseased fish in RAS | C. M. Millard et al., Appl Environ Microbiol 78, 8219-8226 (2012). |
| QMA0246 | R4 | SAMN09104511 | *Streptococcus iniae* | *Lates calcarifer* | 2009 | Australia: SA | Allied Biotechnology Pty LTd | Isolated from diseased fish in RAS | C. M. Millard et al., Appl Environ Microbiol 78, 8219-8226 (2012). |
| QMA0247 | R5 | SAMN09104512 | *Streptococcus iniae* | *Lates calcarifer* | 2009 | Australia: SA | Allied Biotechnology Pty LTd | Isolated from diseased fish in RAS | C. M. Millard et al., Appl Environ Microbiol 78, 8219-8226 (2012). |
| QMA0248 | R6 | SAMN09104513 | *Streptococcus iniae* | *Lates calcarifer* | 2009 | Australia: SA | Allied Biotechnology Pty LTd | Isolated from diseased fish in RAS | C. M. Millard et al., Appl Environ Microbiol 78, 8219-8226 (2012). |
| QMA0249 | R7 | SAMN09104514 | *Streptococcus iniae* | *Lates calcarifer* | 2009 | Australia: SA | Allied Biotechnology Pty LTd | Isolated from diseased fish in RAS | C. M. Millard et al., Appl Environ Microbiol 78, 8219-8226 (2012). |
| QMA0250 | I6 | SAMN09104515 | *Streptococcus iniae* | *Lates calcarifer* | 2007 | Australia: NSW | Allied Biotechnology Pty LTd | included in autogenous vaccine 14/1/08 | C. M. Millard et al., Appl Environ Microbiol 78, 8219-8226 (2012). |
| QMA0251 | I7 | SAMN09104516 | *Streptococcus iniae* | *Lates calcarifer* | 2008 | Australia: NSW | Allied Biotechnology Pty LTd | included in autogenous vaccine 30/7/08 | C. M. Millard et al., Appl Environ Microbiol 78, 8219-8226 (2012). |
| QMA0252 | I8 | SAMN09104517 | *Streptococcus iniae* | *Lates calcarifer* | 2008 | Australia: NSW | Allied Biotechnology Pty LTd | included in autogenous vaccine 30/7/08 | C. M. Millard et al., Appl Environ Microbiol 78, 8219-8226 (2012). |
| QMA0253 | I9 | SAMN09104518 | *Streptococcus iniae* | *Lates calcarifer* | 2009 | Australia: NSW | Allied Biotechnology Pty LTd | included in autogenous vaccine 16/3/09 | C. M. Millard et al., Appl Environ Microbiol 78, 8219-8226 (2012). |
| QMA0254 | I10 | SAMN09104519 | *Streptococcus iniae* | *Lates calcarifer* | 2009 | Australia: NSW | Allied Biotechnology Pty LTd | included in autogenous vaccine 16/3/09 | C. M. Millard et al., Appl Environ Microbiol 78, 8219-8226 (2012). |
| QMA0258 | MP2 | SAMN09104520 | *Streptococcus iniae* | *Lates calcarifer* | 2008 | Australia: QLD | Allied Biotechnology Pty LTd | Isolated from diseased fish in earthen ponds | |
| QMA0371 | 140/11 | SAMN09104521 | *Streptococcus iniae* | *Scortum barcoo* | 2011 | Australia: NSW | Tréidlia Biovet Pty. Ltd | Isolated from diseased fish in RAS | |
| QMA0373 | P14723.6 | SAMN09104522 | *Streptococcus iniae* | *Lates calcarifer* | 2012 | Australia: QLD | Tréidlia Biovet Pty. Ltd | Isolated from diseased fish in earthen ponds | |
| QMA0374 | P14723.8 | SAMN09104523 | *Streptococcus iniae* | *Lates calcarifer* | 2012 | Australia: QLD | Tréidlia Biovet Pty. Ltd | Isolated from diseased fish in earthen ponds | |
| QMA0445 | Si04 | SAMN09104524 | *Streptococcus iniae* | *Oreochromis sp.* | 1998 | USA: Texas | Novartis Animal Health | | |
| QMA0446 | MT2376 | SAMN09104525 | *Streptococcus iniae* | *Oreochromis sp.* | 1998 | USA: Texas | Novartis Animal Health | also in Marine Scotland, Aberdeen collection | A. C. Barnes, F. M. Young, M. T. Horne, A. E. Ellis, *Diseases of Aquatic Organisms* **53**, 241-247 (2003). |
| QMA0447 | MT2374 | SAMN09104526 | *Streptococcus iniae* | *Morone saxatilis x M. chrysops* | 1996 | USA: Maine | Novartis Animal Health | also in Marine Scotland, Aberdeen collection | A. C. Barnes, F. M. Young, M. T. Horne, A. E. Ellis, *Diseases of Aquatic Organisms* **53**, 241-247 (2003). |
| QMA0448 | MT2378 | SAMN09104527 | *Streptococcus iniae* | *Morone saxatilis x M. chrysops* | 1998 | USA: Texas | Novartis Animal Health | also in Marine Scotland, Aberdeen collection | A. C. Barnes, F. M. Young, M. T. Horne, A. E. Ellis, *Diseases of Aquatic Organisms* **53**, 241-247 (2003). |
| QMA0457 | Si18 | SAMN09104528 | *Streptococcus iniae* | *Oreochromis sp.* | 2005 | USA: Idaho | Novartis Animal Health | | |
| QMA0458 | Si20 | SAMN09104529 | *Streptococcus iniae* | *Epalzeorhynchos bicolor* | 2004 | USA: Florida | Novartis Animal Health | | |
| QMA0462 | Si24 | SAMN09104530 | *Streptococcus iniae* | *Chromobotia macracanthus* | 2005 | USA: Florida | Novartis Animal Health | | |
| QMA0463 | Si25 | SAMN09104531 | *Streptococcus iniae* | *Chromobotia macracanthus* | 2005 | USA: Florida | Novartis Animal Health | | |
| QMA0466 | Si28 | SAMN09104532 | *Streptococcus iniae* | *Oreochromis sp.* | missing | USA: Idaho | Novartis Animal Health | | |
| QMA0467 | Si29 | SAMN09104533 | *Streptococcus iniae* | *Epalzeothynchos frenatum* | 2004 | USA: Florida | Novartis Animal Health | | |
| QMA0468 | Si30 | SAMN09104534 | *Streptococcus iniae* | *Oreochromis sp.* | 2005 | USA: Minnesota | Novartis Animal Health | | |
| QMA0490 | SA06 | SAMN09104535 | *Streptococcus iniae* | *Oreochromis sp.* | 2015 | Honduras | Regal Springs Tilapia | | |
| QMA0491 | SA07 | SAMN09104536 | *Streptococcus iniae* | *Oreochromis sp.* | 2015 | Honduras | Regal Springs Tilapia | | |
| QMA0492 | SA08 | SAMN09104537 | *Streptococcus iniae* | *Oreochromis sp.* | 2015 | Honduras | Regal Springs Tilapia | | |
| QMA0493 | SA09 | SAMN09104538 | *Streptococcus iniae* | *Oreochromis sp.* | 2016 | Honduras | Regal Springs Tilapia | | |
