## Supplementary Table 4 for "Evolutionary epidemiology of *Streptococcus iniae*: linking mutation rate dynamics with adaptation to novel immunological landscapes"

| Assembly | Accession Number | # contigs | Largest conti | Total length | GC (%) | N50 | Coverage |
|---|---|---|---|---|---|---|---|
| QMA0071 | QLSO00000000 | 78 | 113807 | 1960490 | 36.54 | 47258 | *70* |
| QMA0074 | QLSN00000000 | 90 | 114060 | 1988879 | 36.55 | 46257 | 75 |
| QMA0077 | QLSM00000000 | 88 | 113957 | 1989678 | 36.54 | 47212 | 68 |
| QMA0078 | QLSL00000000 | 78 | 113807 | 1961453 | 36.54 | 47302 | 63 |
| QMA0080 | QLSK00000000 | 92 | 113816 | 1960466 | 36.58 | 39635 | 99 |
| QMA0082 | QLSJ00000000 | 103 | 113816 | 1973124 | 36.59 | 39635 | 60 |
| QMA0083 | QLSI00000000 | 100 | 145495 | 2036785 | 36.59 | 48872 | 143 |
| QMA0084 | QLSH00000000 | 79 | 244206 | 2017325 | 36.57 | 46107 | 156 |
| QMA0087 | QLSG00000000 | 80 | 145495 | 2000434 | 36.58 | 47143 | 60 |
| QMA0130 | QLSF00000000 | 79 | 145698 | 1997141 | 36.57 | 48787 | 176 |
| QMA0131 | QLSE00000000 | 77 | 146714 | 1995984 | 36.55 | 48787 | 182 |
| QMA0133 | QLSD00000000 | 94 | 206678 | 2013947 | 36.6 | 50804 | 168 |
| QMA0134 | QLSC00000000 | 78 | 206585 | 1998327 | 36.57 | 51251 | 126 |
| QMA0135 | QLSB00000000 | 77 | 206613 | 1996612 | 36.57 | 50185 | 120 |
| QMA0137 | QLSA00000000 | 79 | 206679 | 1998125 | 36.57 | 48674 | 130 |
| QMA0138 | QLRZ00000000 | 76 | 145698 | 1997111 | 36.59 | 50804 | 126 |
| QMA0139 | QLRY00000000 | 88 | 122959 | 2185272 | 36.59 | 44717 | 120 |
| QMA0140 | QLRX00000000 | 80 | 129206 | 2079333 | 36.46 | 44887 | 128 |
| QMA0141 | QLRW00000000 | 69 | 202528 | 2174028 | 36.34 | 64558 | 161 |
| QMA0142 | QLRV00000000 | 79 | 170545 | 1873102 | 36.5 | 36283 | 112 |
| QMA0150 | QLRU00000000 | 77 | 170479 | 1880573 | 36.54 | 42139 | 68 |
| QMA0155 | QLRT00000000 | 79 | 146417 | 1994355 | 36.54 | 51746 | 128 |
| QMA0156 | QLRS00000000 | 81 | 108283 | 1988499 | 36.52 | 48079 | 132 |
| QMA0157 | QLRR00000000 | 83 | 145495 | 1993093 | 36.56 | 47258 | 72 |
| QMA0158 | QLRQ00000000 | 80 | 145495 | 1993353 | 36.57 | 47262 | 91 |
| QMA0159 | QLRP00000000 | 73 | 145495 | 1991061 | 36.53 | 51215 | 132 |
| QMA0160 | QLRO00000000 | 81 | 145495 | 1988650 | 36.52 | 47212 | 121 |
| QMA0161 | QLRN00000000 | 77 | 145495 | 1996299 | 36.56 | 48872 | 70 |
| QMA0162 | QLRM00000000 | 79 | 145495 | 1995292 | 36.57 | 47258 | 70 |
| QMA0163 | QLRL00000000 | 77 | 145495 | 1993641 | 36.57 | 48550 | 75 |
| QMA0164 | QLRK00000000 | 87 | 114062 | 1987989 | 36.54 | 47212 | 72 |
| QMA0165 | QLRJ00000000 | 89 | 114062 | 1988545 | 36.53 | 46669 | 78 |
| QMA0177 | QLRI00000000 | 81 | 170773 | 1879603 | 36.52 | 42139 | 140 |
| QMA0180 | QLRH00000000 | 80 | 169960 | 1884134 | 36.54 | 42622 | 98 |
| QMA0186 | QLRG00000000 | 73 | 210824 | 1903339 | 36.5 | 47015 | 104 |
| QMA0187 | QLRF00000000 | 79 | 165781 | 2044301 | 36.49 | 48254 | 120 |
| QMA0188 | QLRE00000000 | 71 | 154296 | 1896879 | 36.47 | 48529 | 144 |
| QMA0189 | QLRD00000000 | 74 | 211771 | 1883574 | 36.46 | 47016 | 144 |
| QMA0190 | QLRC00000000 | 88 | 111205 | 2030860 | 36.54 | 46236 | 88 |
| QMA0191 | QLRB00000000 | 79 | 170929 | 1882523 | 36.53 | 42139 | 132 |
| QMA0207 | QLRA00000000 | 81 | 142777 | 1878618 | 36.54 | 42139 | 98 |
| QMA0216 | QLQZ00000000 | 78 | 137049 | 1965106 | 36.54 | 47212 | 81 |
| QMA0218 | QLQY00000000 | 87 | 114062 | 1989588 | 36.53 | 47817 | 105 |
| QMA0220 | QLQX00000000 | 81 | 145495 | 1990891 | 36.54 | 47258 | 91 |
| QMA0221 | QLQW00000000 | 77 | 145495 | 1994241 | 36.56 | 47258 | 104 |
| QMA0222 | QLQV00000000 | 78 | 145495 | 1990796 | 36.56 | 47258 | 104 |
| QMA0233 | QLQU00000000 | 96 | 100462 | 2044133 | 36.63 | 37181 | 108 |

| | | | | | | | |
|---|---|---|---|---|---|---|---|
| QMA0234 | QLQT00000000 | 93 | 100734 | 2047858 | 36.64 | 38189 | 60 |
| QMA0235 | QLQS00000000 | 96 | 100879 | 2046848 | 36.63 | 38039 | 60 |
| QMA0236 | QLQR00000000 | 92 | 100000 | 2049063 | 36.65 | 38188 | 135 |
| QMA0244 | QLSP00000000 | 80 | 145495 | 1996827 | 36.58 | 48550 | 70 |
| QMA0245 | QLQQ00000000 | 79 | 145495 | 1997464 | 36.58 | 48427 | 65 |
| QMA0246 | QLQP00000000 | 79 | 145495 | 1997537 | 36.58 | 48550 | 60 |
| QMA0247 | QLQO00000000 | 84 | 145495 | 1992338 | 36.57 | 47258 | 77 |
| QMA0248 | CP022392.1 | 77 | 145495 | 1994521 | 36.56 | 51359 | 117 |
| QMA0249 | QLQN00000000 | 98 | 94661 | 2038042 | 36.64 | 38039 | 65 |
| QMA0250 | QLQM00000000 | 84 | 145495 | 1992393 | 36.52 | 46250 | 130 |
| QMA0251 | QLQL00000000 | 78 | 145495 | 1937737 | 36.52 | 46250 | 77 |
| QMA0252 | QLQK00000000 | 78 | 145495 | 1938644 | 36.52 | 47260 | 70 |
| QMA0253 | QLQJ00000000 | 96 | 100544 | 2047061 | 36.64 | 38047 | 143 |
| QMA0254 | QLQI00000000 | 97 | 100734 | 2046706 | 36.64 | 38047 | 66 |
| QMA0258 | QLQH00000000 | 79 | 137165 | 1957491 | 36.49 | 47212 | 72 |
| QMA0371 | QLQG00000000 | 110 | 113162 | 2211973 | 36.6 | 38190 | 78 |
| QMA0373 | QLQF00000000 | 89 | 114017 | 1990274 | 36.51 | 47729 | 136 |
| QMA0374 | QLQE00000000 | 86 | 114567 | 1988104 | 36.54 | 47729 | 135 |
| QMA0445 | QLQD00000000 | 89 | 167904 | 2264970 | 36.61 | 46597 | 120 |
| QMA0446 | QLQC00000000 | 93 | 167904 | 2263915 | 36.57 | 46597 | 120 |
| QMA0447 | QLQB00000000 | 73 | 206612 | 1960937 | 36.53 | 50804 | 144 |
| QMA0448 | QLQA00000000 | 75 | 206670 | 1994040 | 36.55 | 50183 | 143 |
| QMA0457 | QLPZ00000000 | 87 | 180074 | 2037029 | 36.52 | 47212 | 154 |
| QMA0458 | QLPY00000000 | 80 | 146729 | 1980652 | 36.52 | 48024 | 156 |
| QMA0462 | QLPX00000000 | 87 | 139666 | 1998379 | 36.58 | 45485 | 161 |
| QMA0463 | QLPW00000000 | 100 | 244491 | 1978846 | 36.6 | 44447 | 144 |
| QMA0466 | QLPV00000000 | 82 | 145698 | 1993146 | 36.53 | 46249 | 150 |
| QMA0467 | QLPU00000000 | 75 | 145713 | 1974500 | 36.55 | 48002 | 144 |
| QMA0468 | QLPT00000000 | 90 | 145642 | 2037588 | 36.53 | 47212 | 121 |
| QMA0490 | QLPS00000000 | 78 | 145598 | 1960320 | 36.53 | 46062 | 63 |
| QMA0491 | QLPR00000000 | 82 | 145598 | 1966151 | 36.56 | 46399 | 72 |
| QMA0492 | QLPQ00000000 | 78 | 145598 | 1965383 | 36.56 | 46399 | 70 |
| QMA0493 | QLPP00000000 | 79 | 145598 | 1965971 | 36.56 | 46062 | 77 |
