## Supplementary Table 5 for "Evolutionary epidemiology of *Streptococcus iniae*: linking mutation rate dynamics with adaptation to novel immunological landscapes"

| Node | Clade | Total SNPs | Num of SNPs | Num of SNPs | Num of Reco | Bases in Rec | r/m | rho/theta | Genome Len | Bases in Clo |
|---|---|---|---|---|---|---|---|---|---|---|
| **QMA0248** | A | 8 | 4 | 4 | 1 | 19794 | 1 | 0.25 | 1643310 | 1623580 |
| **QMA0071** | A | 1 | 0 | 1 | 0 | 19728 | 0 | 0 | 1643311 | 1623581 |
| **QMA0078** | A | 3 | 0 | 3 | 0 | 19728 | 0 | 0 | 1643311 | 1623581 |
| **QMA0083** | A | 1 | 0 | 1 | 0 | 19728 | 0 | 0 | 1643310 | 1623580 |
| **QMA0087** | A | 2 | 0 | 2 | 0 | 19728 | 0 | 0 | 1643310 | 1623580 |
| **QMA0155** | A | 0 | 0 | 0 | 0 | 19728 | 0 | 0 | 1643310 | 1623580 |
| **QMA0156** | A | 0 | 0 | 0 | 0 | 19728 | 0 | 0 | 1643310 | 1623580 |
| **QMA0157** | A | 0 | 0 | 0 | 0 | 19728 | 0 | 0 | 1643310 | 1623580 |
| **QMA0158** | A | 9 | 0 | 9 | 0 | 19728 | 0 | 0 | 1643309 | 1623579 |
| **QMA0159** | A | 0 | 0 | 0 | 0 | 19728 | 0 | 0 | 1643310 | 1623580 |
| **QMA0160** | A | 7 | 0 | 7 | 0 | 19728 | 0 | 0 | 1643312 | 1623582 |
| **QMA0161** | A | 2 | 0 | 2 | 0 | 19728 | 0 | 0 | 1643312 | 1623582 |
| **QMA0162** | A | 3 | 0 | 3 | 0 | 19728 | 0 | 0 | 1643312 | 1623582 |
| **QMA0163** | A | 2 | 0 | 2 | 0 | 19728 | 0 | 0 | 1643312 | 1623582 |
| **QMA0216** | A | 6 | 0 | 6 | 0 | 19728 | 0 | 0 | 1643310 | 1623580 |
| **QMA0220** | A | 0 | 0 | 0 | 0 | 19728 | 0 | 0 | 1643310 | 1623580 |
| **QMA0221** | A | 0 | 0 | 0 | 0 | 19728 | 0 | 0 | 1643310 | 1623580 |
| **QMA0222** | A | 0 | 0 | 0 | 0 | 19728 | 0 | 0 | 1643308 | 1623578 |
| **QMA0244** | A | 0 | 0 | 0 | 0 | 19728 | 0 | 0 | 1643312 | 1623582 |
| **QMA0245** | A | 0 | 0 | 0 | 0 | 19728 | 0 | 0 | 1643312 | 1623582 |
| **QMA0246** | A | 0 | 0 | 0 | 0 | 19728 | 0 | 0 | 1643312 | 1623582 |
| **QMA0247** | A | 0 | 0 | 0 | 0 | 19728 | 0 | 0 | 1643310 | 1623580 |
| **QMA0250** | A | 1 | 0 | 1 | 0 | 19728 | 0 | 0 | 1643310 | 1623580 |
| **QMA0251** | A | 0 | 0 | 0 | 0 | 19728 | 0 | 0 | 1643310 | 1623580 |
| **QMA0252** | A | 0 | 0 | 0 | 0 | 19728 | 0 | 0 | 1643310 | 1623580 |
| **QMA0258** | A | 0 | 0 | 0 | 0 | 19728 | 0 | 0 | 1643311 | 1623581 |
| **QMA0371** | A | 81 | 22 | 59 | 1 | 19728 | 0.372881 | 0.016949 | 1643312 | 1623582 |
| **QMA0457** | A | 0 | 0 | 0 | 0 | 19728 | 0 | 0 | 1643309 | 1623579 |
| **QMA0458** | A | 0 | 0 | 0 | 0 | 19728 | 0 | 0 | 1643315 | 1623585 |
| **QMA0467** | A | 2 | 0 | 2 | 0 | 19728 | 0 | 0 | 1643315 | 1623585 |
| **QMA0468** | A | 19 | 0 | 19 | 0 | 19728 | 0 | 0 | 1643304 | 1623574 |
| **QMA0490** | A | 0 | 0 | 0 | 0 | 19728 | 0 | 0 | 1643319 | 1623589 |
| **QMA0491** | A | 0 | 0 | 0 | 0 | 19728 | 0 | 0 | 1643319 | 1623589 |
| **QMA0492** | A | 0 | 0 | 0 | 0 | 19728 | 0 | 0 | 1643319 | 1623589 |
| **QMA0493** | A | 0 | 0 | 0 | 0 | 19728 | 0 | 0 | 1643319 | 1623589 |
| **QMA0084** | B | 17 | 0 | 17 | 0 | 19728 | 0 | 0 | 1643309 | 1623579 |
| **QMA0462** | B | 1 | 0 | 1 | 0 | 19728 | 0 | 0 | 1643309 | 1623579 |
| **QMA0463** | B | 0 | 0 | 0 | 0 | 19728 | 0 | 0 | 1643309 | 1623579 |
| **QMA0080** | C1 | 3 | 0 | 3 | 0 | 19728 | 0 | 0 | 1643295 | 1623565 |
| **QMA0082** | C1 | 0 | 0 | 0 | 0 | 19728 | 0 | 0 | 1643296 | 1623566 |
| **QMA0142** | C1 | 1 | 0 | 1 | 0 | 19728 | 0 | 0 | 1643292 | 1623562 |
| **QMA0150** | C1 | 1 | 0 | 1 | 0 | 19728 | 0 | 0 | 1643292 | 1623562 |
| **QMA0177** | C1 | 1 | 0 | 1 | 0 | 19728 | 0 | 0 | 1643292 | 1623562 |
| **QMA0180** | C1 | 0 | 0 | 0 | 0 | 19728 | 0 | 0 | 1643292 | 1623562 |
| **QMA0191** | C1 | 0 | 0 | 0 | 0 | 19728 | 0 | 0 | 1643292 | 1623562 |
| **QMA0207** | C1 | 0 | 0 | 0 | 0 | 19728 | 0 | 0 | 1643292 | 1623562 |
| **QMA0074** | C2 | 31 | 6 | 25 | 1 | 19728 | 0.24 | 0.04 | 1643307 | 1623577 |
| **QMA0077** | C2 | 91 | 22 | 69 | 1 | 19728 | 0.318841 | 0.014493 | 1643310 | 1623580 |
| **QMA0164** | C2 | 0 | 0 | 0 | 0 | 19728 | 0 | 0 | 1643311 | 1623581 |
| **QMA0165** | C2 | 0 | 0 | 0 | 0 | 19728 | 0 | 0 | 1643311 | 1623581 |
| **QMA0218** | C2 | 0 | 0 | 0 | 0 | 19728 | 0 | 0 | 1643311 | 1623581 |
| **QMA0373** | C2 | 3 | 0 | 3 | 0 | 19728 | 0 | 0 | 1643309 | 1623579 |
| **QMA0374** | C2 | 0 | 0 | 0 | 0 | 19728 | 0 | 0 | 1643309 | 1623579 |
| **QMA0186** | D | 0 | 0 | 0 | 0 | 19728 | 0 | 0 | 1643313 | 1623583 |
| **QMA0188** | D | 16 | 0 | 16 | 0 | 19728 | 0 | 0 | 1643311 | 1623581 |
| **QMA0189** | D | 0 | 0 | 0 | 0 | 19728 | 0 | 0 | 1643311 | 1623581 |
| **QMA0133** | E1 | 6 | 0 | 6 | 0 | 19728 | 0 | 0 | 1643318 | 1623588 |
| **QMA0134** | E1 | 0 | 0 | 0 | 0 | 19728 | 0 | 0 | 1643318 | 1623588 |
| **QMA0135** | E1 | 0 | 0 | 0 | 0 | 19728 | 0 | 0 | 1643318 | 1623588 |
| **QMA0137** | E1 | 12 | 0 | 12 | 0 | 19728 | 0 | 0 | 1643318 | 1623588 |
| **QMA0138** | E1 | 15 | 0 | 15 | 0 | 19728 | 0 | 0 | 1643317 | 1623587 |
| **QMA0447** | E1 | 0 | 0 | 0 | 0 | 19728 | 0 | 0 | 1643319 | 1623589 |
| **QMA0448** | E1 | 3 | 0 | 3 | 0 | 19728 | 0 | 0 | 1643319 | 1623589 |
| **QMA0130** | E2 | 13 | 0 | 13 | 0 | 19728 | 0 | 0 | 1643465 | 1623735 |
| **QMA0131** | E2 | 0 | 0 | 0 | 0 | 19728 | 0 | 0 | 1643465 | 1623735 |
| **QMA0466** | E2 | 48 | 0 | 48 | 0 | 19728 | 0 | 0 | 1643318 | 1623588 |
| **QMA0139** | F | 84 | 0 | 84 | 0 | 19616 | 0 | 0 | 1643312 | 1623695 |
| **QMA0190** | F | 0 | 0 | 0 | 0 | 19616 | 0 | 0 | 1643311 | 1623694 |
| **QMA0233** | F | 3 | 0 | 3 | 0 | 19615 | 0 | 0 | 1643309 | 1623692 |

| | | | | | | | | | | |
|---|---|---|---|---|---|---|---|---|---|---|
| **QMA0234** | F | 0 | 0 | 0 | 0 | 19615 | 0 | 0 | 1643309 | 1623692 |
| **QMA0235** | F | 1 | 0 | 1 | 0 | 19615 | 0 | 0 | 1643309 | 1623692 |
| **QMA0236** | F | 2 | 0 | 2 | 0 | 19615 | 0 | 0 | 1643309 | 1623692 |
| **QMA0249** | F | 10 | 0 | 10 | 0 | 19615 | 0 | 0 | 1643308 | 1623691 |
| **QMA0253** | F | 1 | 0 | 1 | 0 | 19615 | 0 | 0 | 1643309 | 1623692 |
| **QMA0254** | F | 1 | 0 | 1 | 0 | 19615 | 0 | 0 | 1643309 | 1623692 |
| **QMA0445** | G | 9 | 0 | 9 | 0 | 19616 | 0 | 0 | 1643305 | 1623688 |
| **QMA0446** | G | 0 | 0 | 0 | 0 | 19616 | 0 | 0 | 1643310 | 1623693 |
| **QMA0140** | U | 0 | 0 | 0 | 0 | 19324 | 0 | 0 | 1643317 | 1623992 |
| **QMA0141** | U | 13752 | 5428 | 8324 | 95 | 126301 | 0.65209 | 0.011413 | 1643282 | 1623957 |
| **QMA0187** | U | 192 | 0 | 192 | 0 | 19616 | 0 | 0 | 1643311 | 1623694 |
